# Two populations with different cell dynamics maintain urethral epithelial homeostasis in female and male mice

**DOI:** 10.64898/2026.10.16.759572

**Authors:** Ioanna Kaklamanou, Uday Ram Gubbala, Oscar Bay Axelsen, Diana Andrejeva, Edouard Hannezo, Mariaceleste Aragona

## Abstract

The urethra is an understudied organ in both humans and animal models. Here we investigate the mouse urethra to gain insights into its tissue architecture, maintenance mechanisms, and epithelial cell dynamics in homeostasis. For this, we generated a single-cell RNA sequencing dataset that includes cells from the entire urethra of both female and male, with detailed regional information. Using lineage tracing and cell proliferation assays, we focused on two potential progenitor cell types, the Keratin 5^+^ and Keratin 14^+^ cells, which we found both capable of generating the full thickness of the epithelium, yet with opposite and complementary patterns of self-renewal. Our results further highlight key differences between females and males, as we observed a higher epithelium stratification, a unique pattern of keratins expression and a higher tissue turnover in females compared to males. This work provides valuable insights that can support future tissue engineering efforts aimed at developing urethral grafts for treating urethral disorders.

## Introduction

Every day, a healthy individual urinates 2-10 times on average^1–3^. This frequent exposure to varying volumes of urine necessitates that the organs of the lower urinary tract – the bladder and the urethra – possess a specialized epithelium capable of adapting to rapid changes in lumen volume and replacing cells lost during urination. This epithelium must be tightly regulated, continuously generating new, identical, and differentiated cells to maintain a protective barrier without disrupting its tissue architecture. While many studies have investigated the mouse bladder epithelium to understand how tissue architecture is maintained and which cell populations contribute to the formation of different epithelial layers, comparatively little is known about the urethral epithelium. This lack of research is particularly striking given that one of the most common congenital abnormalities in humans, called hypospadias, directly affects the urethra. Hypospadias affects 1:250 newborn boys on average and begins during gestation with an abnormal development of the urethra, which can lead to the incomplete closure of the urethral opening, incomplete foreskin formation, downward bending of the penis and a more proximal location of the urethral opening^4,5^. Children with severe cases of hypospadias require multiple reconstructive surgeries to reconstruct the urethral epithelium. Since there is a general lack of tissue, the current gold standard therapeutic approach is to generate tissue grafts from the buccal mucosa or distal epithelium from the surrounding skin^6–8^. However, this treatment can lead to adverse effects, such as fistulas, diverticula, strictures, meatal stenosis, increased bending and undesired hair growth^6,8–10^. Due to the tissue deficiency characteristic of hypospadias, any autologous graft of urethral origin will likely need to be bioengineered in the laboratory. The successful production and application of such a graft depend on a thorough understanding of the urethral tissue under normal, homeostatic conditions.

However, very little is known about the architecture of the urethral epithelium, how its morphology changes across the length of the organ, the mechanisms for tissue homeostasis or any sex-specific differences. The majority of the studies on the urethral epithelium have focused on the development of the organ or the region directly connected to the bladder. There, the epithelium is described as transitional, a type of stratified epithelium, with basal, intermediate and luminal/superficial cells. The basal layer is characterized by high expression of Keratin 14 (Krt14) while Keratin 5 (Krt5) is expressed in the basal and intermediate layer^11–13^. The presence of Krt14⁺ and/or Krt5⁺ basal cells as stem or progenitor cells in various epithelial tissues from other organs^14–23^ highlights the importance of investigating urethral basal cells as a potential source of progenitors during tissue homeostasis. However, this has not been investigated. It is also currently unknown if the proximal urethral epithelium architecture is consistent along the whole length of the organ and between sexes, what is the homeostatic cell behavior, what are the markers describing the different urethral cell types and which cells are maintaining the tissue under homeostasis.

Here, we adopt a multiscale approach to define the molecular profiling, single-cell clonal dynamics and whole adult organ architecture, across different organ regions and for both male and female. In particular, we focus on two specific cell populations: the basal Krt14^+^ and the basal-intermediate Krt5^+^ population. We describe their potential to generate the full epithelial thickness using long-term lineage tracing in multiple mouse models. Together, these findings provide a better understanding of the urethra under homeostatic conditions, which may inform future strategies for developing urethral grafts to treat urethral disorders in human.

## Results

### Tissue architecture morphological transitions in the urethral epithelium

To begin characterizing regional differences in urethral architecture between sexes and along the proximal-distal axis, we analyzed Hematoxylin and Eosin (H&E)-stained cross-sections.

The female urethra was divided into proximal and distal regions (Figure 1A). The distal urethra is surrounded by Skene’s glands (Figure 1B, blue), and epithelial morphology changes from a horseshoe shape proximally to a more oval shape distally (Figure 1B). In males, the proximal urethra includes the prostatic and membranous regions, whereas the penile urethra is distal (Figure 1C). Urethral glands are visible in the proximal male urethra (Figure 1D, blue). The female urethral epithelium is more stratified than the male epithelium, with 2-6 versus 1-5 cell layers across regions (Figure 1E-G). This is reflected by a greater epithelial thickness in females (38 μm proximal, 26 μm distal) than in males (15 μm proximal, 16 μm distal) (Figure 1H).

**Figure 1:**
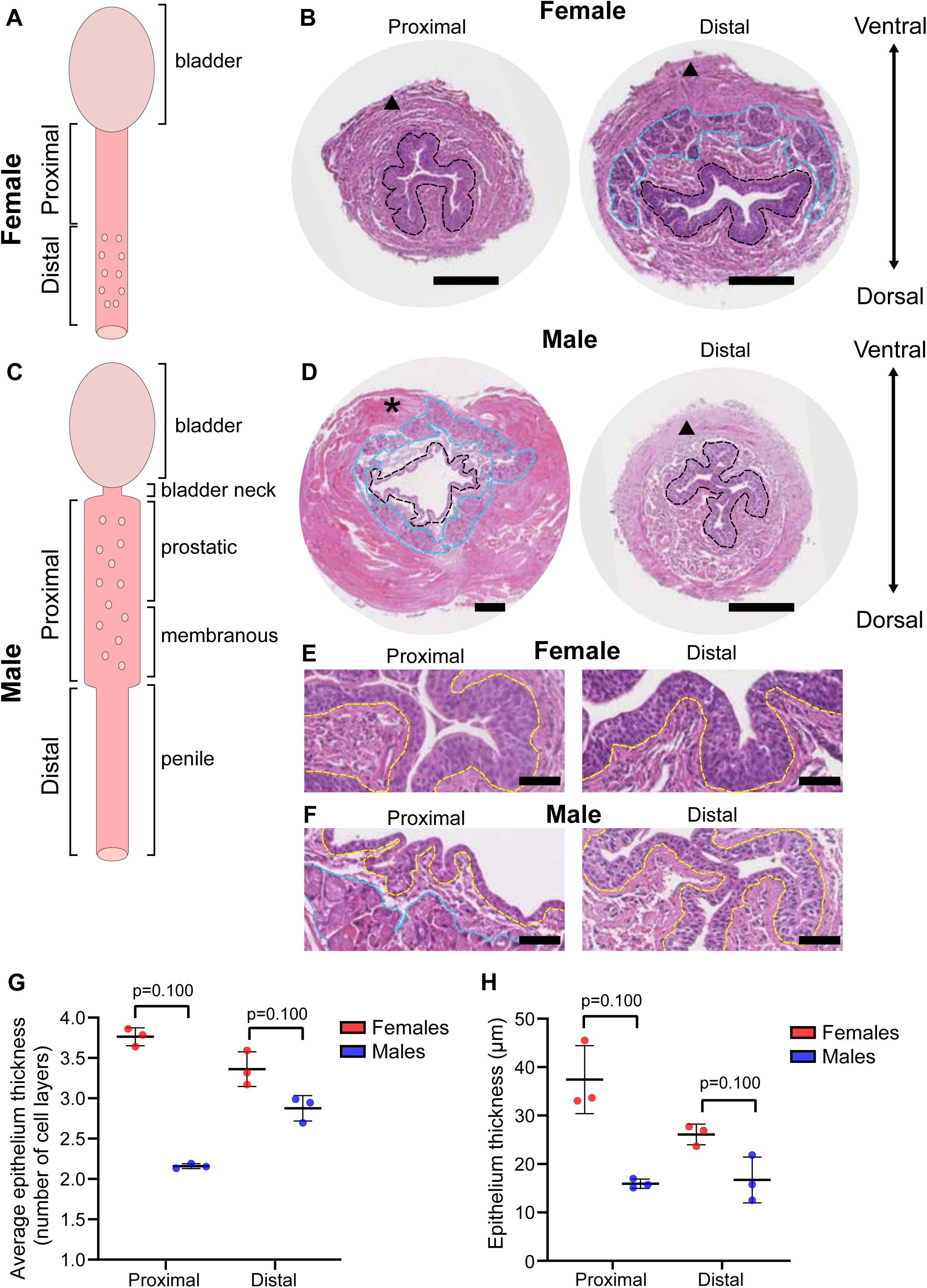
The female and male urethra show gross anatomical and structural differences. A. Schematic of mouse female urethra, the first half being marked as proximal urethra and the second half as distal urethra. The Skene’s glands are visible in the distal urethra. B. H&E staining of the female urethra showing that the morphology of the urethral epithelium (outlined in black) changes from horseshoe shape in proximal urethra to a more elongated oval shape in the distal urethra where the glands (outlined in blue) are visible. Smooth muscle indicated with a triangle. Ventral and dorsal directions indicated with an arrow. Scale bar: 200μm. C. Schematic of mouse male urethra, the first half being marked as proximal (prostatic and membranous) urethra and the second half (penile) as distal urethra. Between bladder and proximal urethra, the circumference of the epithelium is smaller, and it is termed bladder neck. The urethral glands are visible in the proximal urethra. D. H&E staining of the male urethra showing that the morphology of the urethral epithelium (outlined in black) changes from a round shape with thick muscle layer and multiple glands (outlined in blue) in proximal urethra to a more complex shape in the distal urethra, similar to the female epithelium. Striated muscle indicated with an asterisk and smooth muscle indicated with a triangle. Ventral and dorsal directions indicated with an arrow. Scale bar: 200μm E. H&E staining of the female urethra in proximal and distal region. Basement membrane marked with yellow dashed line. Scale bar: 50μm F. H&E staining of the male urethra in proximal and distal region. Basement membrane marked with yellow dashed line and glands outlined with blue. Scale bar: 50μm G. Quantification of the number of cell layers of epithelium in females (red) and males (blue), proximal and distal urethra, n=3 mice per sex. Data represented as mean ± SD. Statistical test is Mann-Whitney U test. H. Quantification of the epithelium thickness in females (red) and males (blue), proximal and distal urethra, n=3 mice per sex. Data represented as mean ± SD. Statistical test is Mann-Whitney U test.

To define where epithelial transitions occur relative to larger tissue changes, we labeled epithelial cells expressing Krt5, a marker of basal and intermediate proliferative cells^12^. Using *Krt5Cre-ERT2;Rosa26^mTmG^*mice (Figure 2A), we visualized cell membranes throughout the urethral epithelium. A high induction dose was used to recombine most epithelial cells, and tissues were analyzed after 24 weeks to allow epithelial turnover. Intact urethras were rendered optically transparent and imaged by light-sheet microscopy (Figure 2B), enabling three-dimensional (3D) reconstruction.

**Figure 2:**
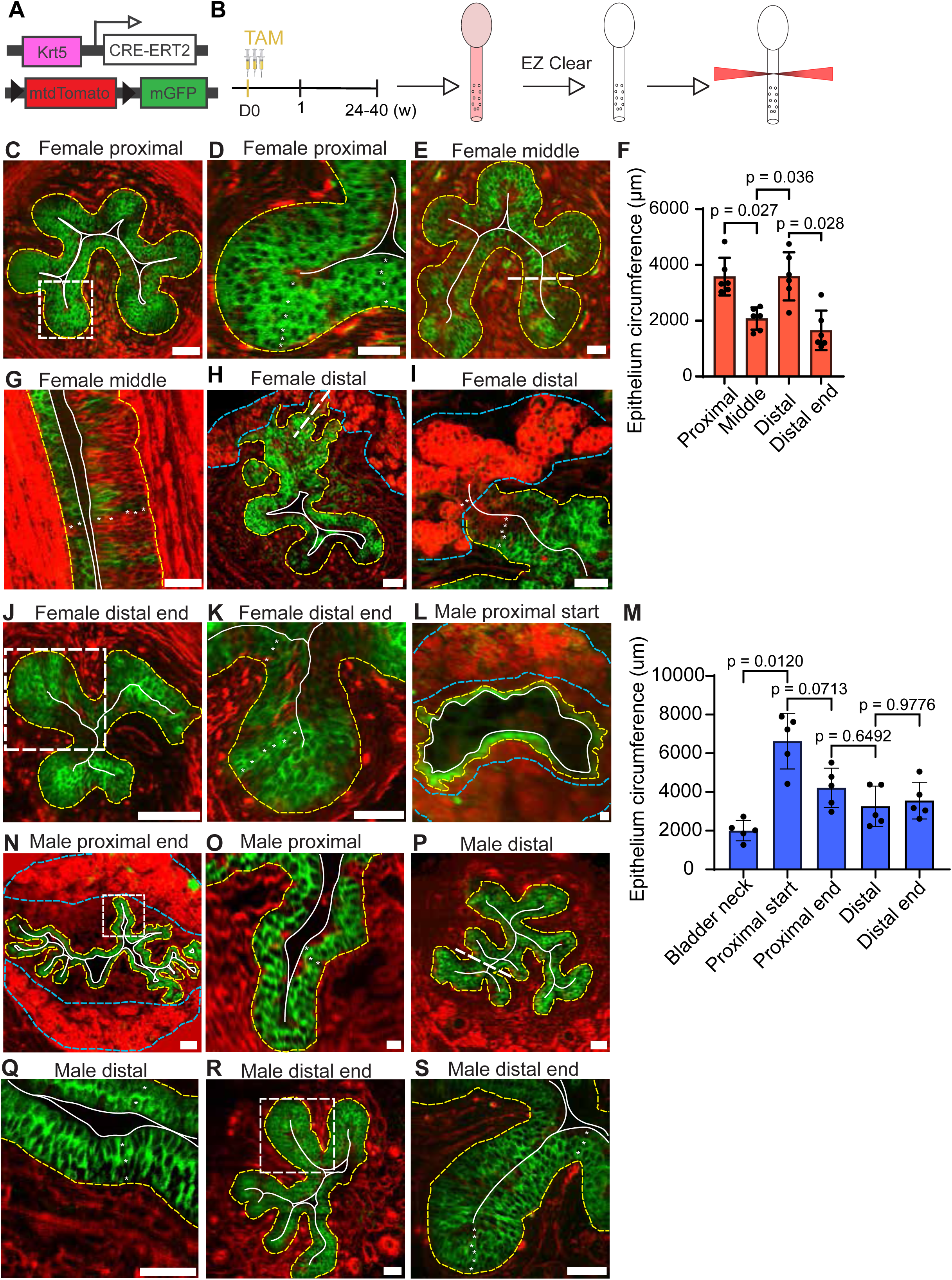
The female and male urethra show tissue architecture differences. A. *Krt5Cre-ERT2*;*Rosa26^mTmG^* mouse construct. B. Experimental strategy. *Krt5Cre-ERT2*;*Rosa26^mTmG^* mice (n=6 females and 5 males) received 3 doses of tamoxifen followed by a 24-40 week chase period. Urethras were then cleared using EZ Clear and imaged on a light sheet microscope. C. Cross-section of female proximal urethra of a *Krt5Cre-ERT2*;*Rosa26^mTmG^* mouse, chased for 26 weeks. White square indicates representative location of Figure 2D. Scale bar: 100μm. D. Epithelium of the female proximal urethra of a *Krt5Cre-ERT2*;*Rosa26^mTmG^* mouse, chased for 26 weeks. Scale bar: 50μm. E. Cross-section of female urethra at the middle region of a *Krt5Cre-ERT2*;*Rosa26^mTmG^* mouse, chased for 26 weeks. White dashed line indicates longitudinal representative location of Figure 2G. Scale bar: 100μm. F. Quantification of the female epithelium circumference at the proximal, middle, distal and distal end regions, n=6 mice. Data represented as mean ± SD. Statistical test is Repeated Measures one-way ANOVA with Geisser-Greenhouse correction and Tukey’s multiple comparisons tests. G. Epithelium of the female urethra of a *Krt5Cre-ERT2*;*Rosa26^mTmG^*mouse at the middle region, chased for 26 weeks. Scale bar: 50μmH. Cross-section of a *Krt5Cre-ERT2*;*Rosa26^mTmG^*distal female urethra, chased for 26 weeks, showing ventrolateral branching of the Skene’s glands’ ducts into the urethral epithelium. White dashed line indicates longitudinal representative location of Figure 2I. Scale bar: 100μm. I. Close-up of duct of Skene’s glands connecting to the urethral epithelium in a *Krt5Cre-ERT2*;*Rosa26^mTmG^* female chased for 26 weeks. Scale bar: 50μm. J. Cross-section of the female urethra of a *Krt5Cre-ERT2*;*Rosa26^mTmG^* mouse at the distal end region, chased for 26 weeks. The closed lumen is outlined with a white line. White square indicates representative location of Figure 2I. Scale bar: 50μm. K. Epithelium of the female urethra of a *Krt5Cre-ERT2*;*Rosa26^mTmG^* mouse at the distal end region, chased for 26 weeks. Scale bar: 50μm. L. Cross-section of male most proximal urethra after the intersection with the reproductive gland ducts at the widened lumen, of a *Krt5Cre-ERT2*;*Rosa26^mTmG^*mouse. Chased for 40 weeks. Scale bar: 100μm.v M. Quantification of the male epithelium circumference at the start and end of bladder neck, proximal and distal urethra, n=5 mice. Data represented as mean ± SD. Statistical test is Repeated Measures one-way ANOVA with Geisser-Greenhouse correction and Tukey’s multiple comparisons tests. N. Cross-section of the middle of the male proximal urethra of a *Krt5Cre-ERT2*;*Rosa26^mTmG^*mouse, surrounded by urethral glands. White square indicates representative location of Figure 2P. Chased for 40 weeks. Scale bar: 100μm. O. Epithelium of the male proximal urethra of a *Krt5Cre-ERT2*;*Rosa26^mTmG^* mouse, chased for 40 weeks. Scale bar: 50μm. P. Cross-section of the male distal urethra of a *Krt5Cre-ERT2*;*Rosa26^mTmG^* mouse, chased for 40 weeks. White dashed line indicates representative longitudinal location of Figure 2R. Scale bar: 100μm. Q. Epithelium of the male distal urethra of a *Krt5Cre-ERT2*;*Rosa26^mTmG^* mouse, chased for 40 weeks. Scale bar: 50μm. R. Cross-section of the male urethra of a *Krt5Cre-ERT2*;*Rosa26^mTmG^* mouse at the distal end, chased for 40 weeks. White square indicates representative location of Figure 2T. Scale bar: 100μm. S. Epithelium of the male urethra of a *Krt5Cre-ERT2*;*Rosa26^mTmG^*mouse at the distal end, chased for 40 weeks. Scale bar: 50μm C-E, G-I, J-L, N-T. Recombined *Krt5Cre-ERT2*;*Rosa26^mTmG^* urethras. Membrane GFP in green, membrane tdTomato in red. Ventral direction is the top of each cross section while dorsal direction is the bottom of each cross section. Basement membranes are marked in yellow, and glands marked in blue. White line indicates apical surface.

In females, the proximal urethra is highly folded (Figure 2C, S1B) and lined by a 4-6-layer transitional epithelium (Figure 2D). In the mid-urethra, folding decreases (Figure S1B), the circumference narrows (Figure 2E,F), and the epithelium adopts cuboidal and columnar morphologies (Figure 2G). Distally, the circumference expands (Figure 2F) as Skene’s gland ducts merge with the epithelial folds (Figure S1C) and cover the ventrolateral urethra (Figure 2H, S1A), where cuboidal urethral epithelium and columnar ductal epithelium coexist (Figure 2I). Near the meatus, the ducts regress and the circumference decreases again (Figure 2F,J), resulting in a stratified cuboidal epithelium (Figure 2K). The lumen reaches its smallest size at the distal end (Figure S1D). Consequently, the female urethra displays three major architectural transitions marked by changes in circumference, epithelial composition, and the presence of Skene’s gland ducts.

In males, the proximal urethra is widest (Figure 2L, 2N) and receives prostatic ducts along its length, while the lumen progressively narrows (Figure 2O, S1F-I). More distally, it becomes highly folded and is lined by a 2-3-layer cuboidal epithelium associated with periurethral glands (Figure 2O). At the membranous urethra, periurethral glands regress and Cowper’s gland ducts appear (Figure S1I), coinciding with a change in lumen shape (Figure 2P) and the appearance of columnar cells in the folds and cuboidal cells centrally (Figure 2Q). Near the glans penis, the urethral shape changes again (Figure 2R) and epithelial stratification increases to 3-5 layers (Figure 2S). Thus, the male urethra undergoes two major architectural transitions: one at the membranous urethra and a second near the glans penis.

### Regional differences in keratins expression

To better characterize epithelial transitions, we examined whether the expression of known keratins varies along the proximal-distal axis and between sexes. Longitudinal sections and whole mounts from both sexes were stained for Krt5, Krt14, Krt8, Krt19, and Krt13 (Figure 3A). Krt5 was uniformly expressed throughout the epithelium in basal and intermediate cells, whereas Krt8 and Krt19 were uniformly expressed in luminal cells and weakly in the intermediate layer. The basal marker Krt14 was detected along the entire female and male urethra but showed a less uniform distribution in the distal male epithelium. Krt13 was present in large patchy areas of the female epithelium, whereas in males it was restricted to small scattered cell clusters. When present, Krt13 was expressed across all three epithelial layers (Figure 3A). Krt20 was detected only in the most proximal female urethra adjacent to the bladder and was absent from males (Figure 3B).

**Figure 3:**
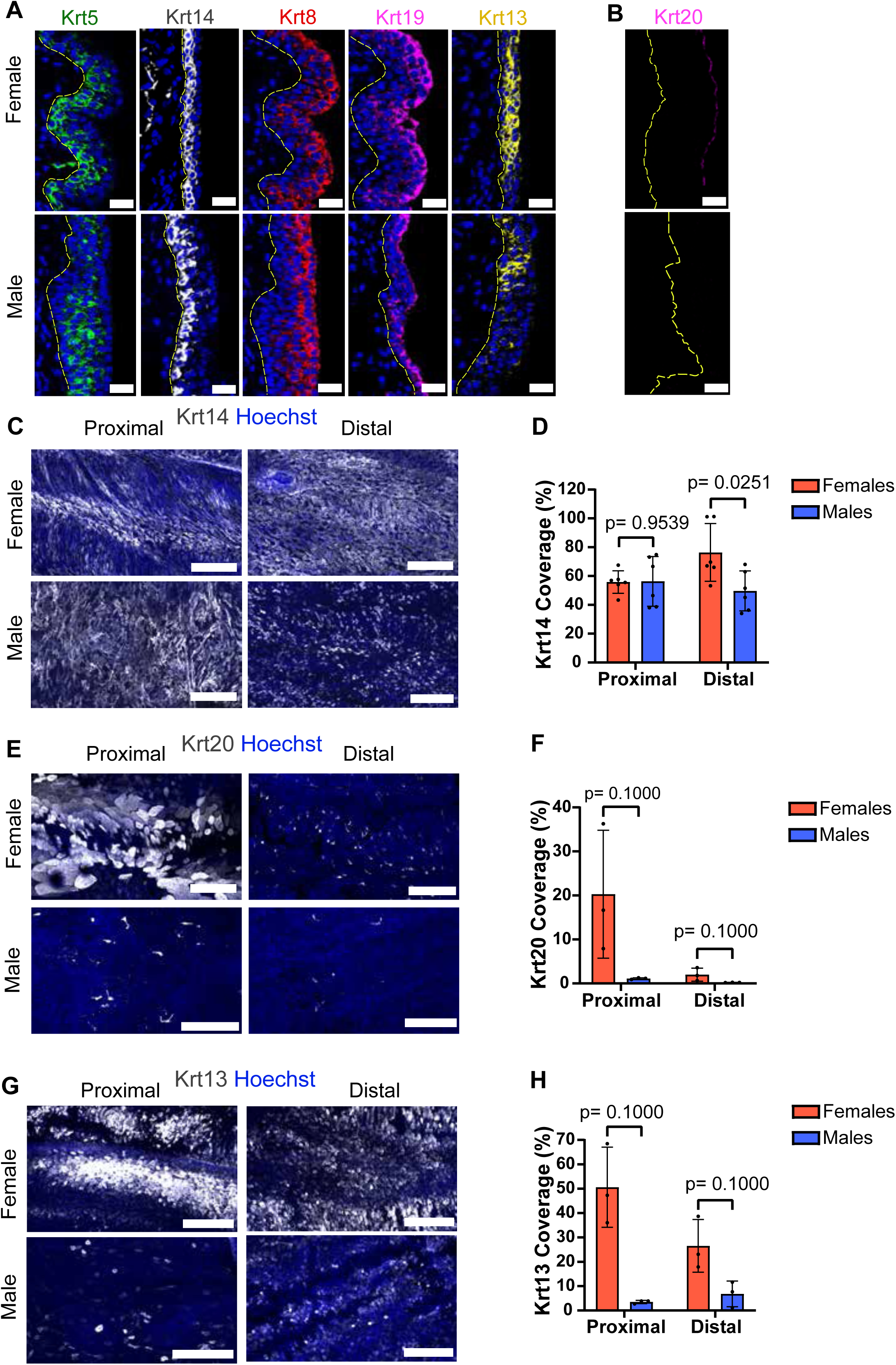
A combination of different keratins can be used to distinguish all the urethral cell layers. A. Immunofluorescence stainings on representative longitudinal sections along the proximal-distal axis for Krt5 (green), Krt14 (white), Krt8 (red), Krt19 (magenta) and Krt13 (yellow). B. Immunofluorescence staining on a longitudinal section for Krt20 (magenta), in a female proximal urethra. A, B. Basement membrane marked with yellow dashed line. Scale bar: 30μm.C. C. Wholemount immunofluorescence staining on a proximal and a distal female and male urethra for Krt14 (gray) and nuclei (Hoechst). Scale bar: 200μm. D. Average coverage of tile scans for Krt14 of female (red) and male (blue) proximal and distal urethras, n=6 mice per sex. Data represented as mean ± SD. Statistical test is unpaired t test with Welch’s correction. E. Wholemount immunofluorescence staining on a proximal and a distal female and male urethra for Krt20 (gray) and nuclei (Hoechst). Scale bar: 200μm. F. Average coverage of tile scans for Krt20 of female (red) and male (blue) proximal and distal urethras, n=3 mice per sex. Data represented as mean ± SD. Statistical test is Mann-Whitney U test. G. Wholemount immunofluorescence staining on a proximal and a distal female and male urethra for Krt13 (gray) and nuclei (Hoechst). Scale bar: 200μm. H. Average coverage of tile scans for Krt13 of female (red) and male (blue) proximal and distal urethras, n=3 mice per sex. Data represented as mean ± SD. Statistical test is Mann-Whitney U test.

Because Krt14, Krt20, and Krt13 showed regional and sex-specific differences, we examined their distribution in flattened whole mounts viewed from the luminal surface. In females, Krt14-expressing cells covered approximately 60% of the proximal region and 80% of the distal region, whereas in males they covered 60% and 50%, respectively (Figure 3E, F). Krt20-expressing cells occupied approximately 20% of the proximal female urethra and resembled bladder umbrella cells, but only 2% of the distal region. In the distal female urethra and in males, Krt20 was instead expressed by small cells with short projections resembling cutaneous neuroendocrine Merkel cells, which are known to express Krt20^31,32^ (Figure 3H). In males, these Krt20-positive neuroendocrine-like cells covered approximately 1% of the proximal region and 0.2% of the distal region (Figure 3E, F).

Comparison of keratin expression with tissue architecture in females showed that the uniform Krt20 domain ended at the first tissue transition, where the epithelium adopted a columnar morphology (Figure S1A, S2A). Krt13-expressing cells covered approximately 50% of the proximal region and 30% of the distal region in females, compared with only 3.5% and 7%, respectively, in males (Figure 3G, H). In females, Krt13 expression was largely continuous along the dorsal side, whereas in males it was sparse, patchy, and irregularly distributed (Figure 3G).

In summary, most basal cells are defined by co-expression of Krt5 and Krt14, intermediate cells by Krt5, Krt8, and Krt19, and luminal cells by region-specific combinations of Krt8, Krt20, Krt19, and, in areas with only two epithelial layers, Krt5 (Figure S2A). Overall, Krt13 and Krt20 show the most pronounced sex-specific differences, whereas Krt14 exhibits more limited variation between males and females.

### Regionally resolved scRNAseq of female and male urethra

To understand if the morphological transitions are reflected in gene expression changes, we performed single-cell RNA sequencing (scRNAseq). We divided the female urethras into proximal and distal and the male urethras in 4 pieces, 2 in proximal and 2 in distal. Each region of each mouse was uniquely barcoded using hashtag antibodies to spatially resolve cells coming from these regions when all the samples are pooled together (Table S1). Live urethra cells were isolated by fluorescence-activated cell sorting (FACS). We sequenced a total of 6419 cells from the female organs from 3 different animals and 5499 cells from the male organs from 3 different animals, of which 3542 female cells and 3765 male cells were bound by a single hashtag antibody (Figure S2B-C, Table S1). In the entire atlas, we uncovered all the different cell types constituting the mouse urethra (Figure 4A, Table S1) with similar proportions between sexes (Figure S2D, Table S1).

**Figure 4:**
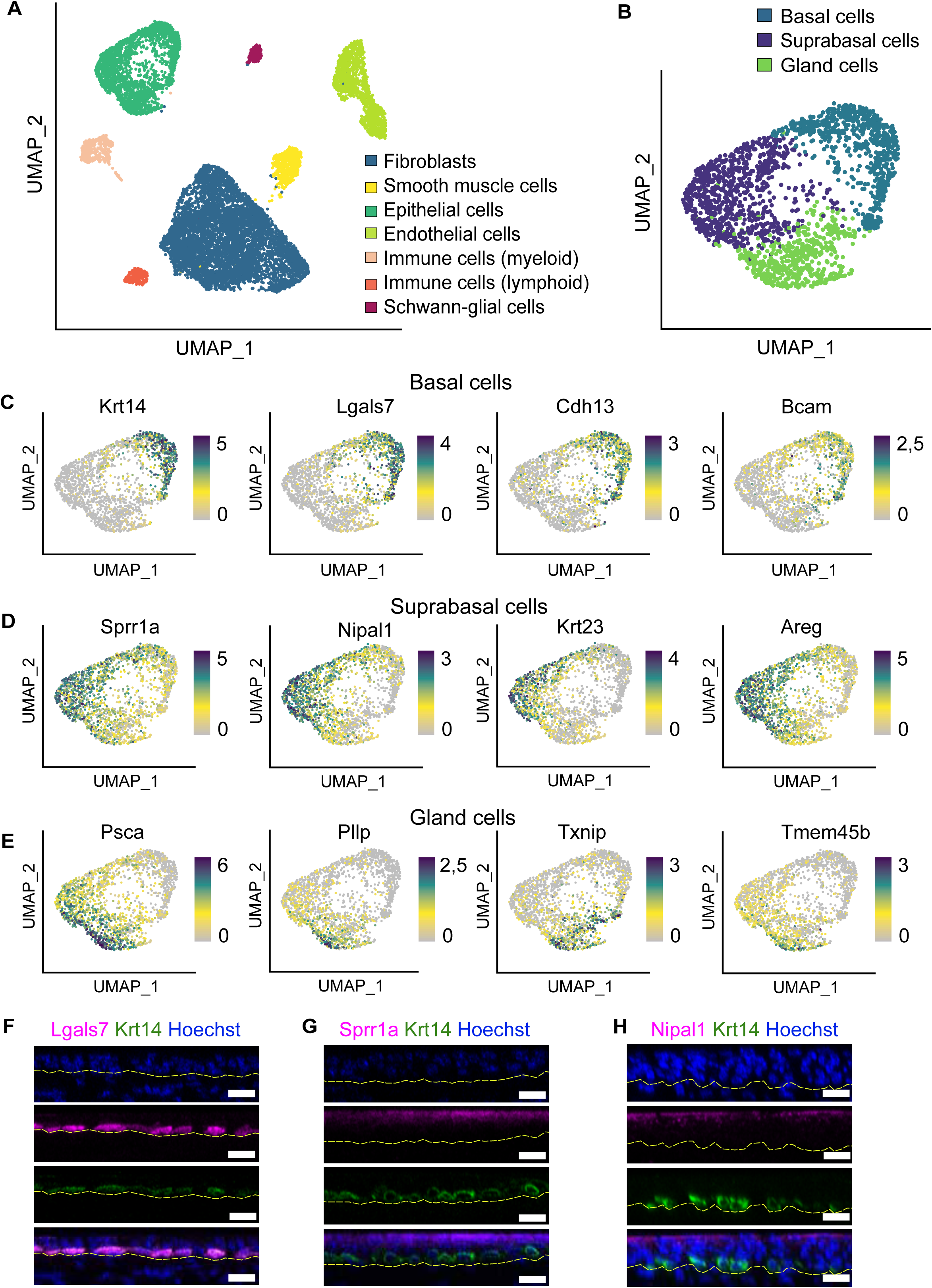
single-cell RNA sequencing identifies marker genes for the urethral epithelium. A. Uniform Manifold Approximation and Projection (UMAP) dimensional reduction of all analyzed urethral cells, n=11918 cells. Seven clusters have been identified. Fibroblasts are in blue, epithelial cells in dark green, endothelial cells in light green, smooth muscle cells in yellow, immune cells of myeloid lineage in beige, immune cells of lymphoid lineage in orange and Schwann-glial cells in magenta. B. UMAP dimensional reduction of all epithelial cells, n=2168 cells. Three clusters have been identified. Basal cells are in blue, suprabasal cells in violet and gland cells in green. C-E. UMAP plot colored by normalized gene expression values for the indicated gene and in the indicated epithelial cluster. Gene expression is visualized as a color gradient going from gray to violet with gray as indicator of no expression and violet as indicator of maximal expression. F. Orthogonal view of wholemount immunofluorescence staining for the basal marker Lgals7 (magenta), Krt14 (green) and nuclei (Hoechst). G. Orthogonal view of wholemount immunofluorescence staining for the luminal marker Sprr1a (magenta), Krt14 (green) and nuclei (Hoechst). H. Orthogonal view of wholemount immunofluorescence staining for the luminal marker Nipal1 (magenta), Krt14 (green) and nuclei (Hoechst). F-H. Basement membrane marked with yellow dashed line. Scale bar: 20μm.

Given our interest in the identification of gene expression changes between different epithelial regions, we decided to investigate this cluster further (Figure 4B, Table S2). In total, there were 2168 epithelial cells in our dataset. Using graph-based clustering, we identified three different subpopulations based on marker gene expression which could be defined as basal cells, suprabasal cells (intermediate and luminal cells) and cells that expressed genes of glandular epithelium (Figure 4B-E). Basal cells express high levels of *Krt14,* Galectin-7 (*Lgals7),* Cadherin-13 (*Cdh13*) and Basal cell adhesion molecule (*Bcam*) (Figure 4C). Suprabasal cells express high levels of Small proline rich protein 1a (*Sprr1a*), Magnesium transporter NIPA-like domain containing 1 (*Nipal1*), Keratin 23 (*Krt23*) and Amphiregulin *(Areg)* (Figure 4D). Glandular cells express high levels of Prostate stem cell antigen (*Psca*), Plasmolipin (*Pllp*), Thioredoxin interacting protein (*Txnip*) and Transmembrane protein 45b (*Tmem45b*) (Figure 4E). We validated by immunofluorescence the expression of the newly identified *Lgals7* as basal marker, *Psca* as glandular marker and *Sprr1a* and *Nipal1* as luminal markers (Figure 4F-H, Figure S2E). The three epithelial clusters had cells belonging to all examined donors (Figure S2F) and regions (Figure S2G). Additionally, the prediction of the cell cycle showed that epithelial cells in G1, G2/M or S phase were homogeneously distributed across each cluster (Figure S2H). When looking at the expression of keratins (Figure S2I) examined before, we found that they were expressed in the expected cell clusters as shown in Figure 3. When comparing cells coming from different regions (proximal or distal) of the same sex, the majority of the Differentially Expressed Genes (DEGs) were found in non-epithelial clusters (Table S3). However, when comparing cells coming from different sexes, the majority of the DEGs were found in the epithelial clusters (Table S3). In total, 190 genes were different between female and male epithelial cells, including known sex chromosome-related genes such as X-inactive specific transcript (*Xist*), Ubiquitous transcriptional regulator Y (*Uty*), DEAD-box helicase 3 Y-linked (*Ddx3y*), Lysine demethylase 5d (*Kdm5d*) and Eukaryotic translation initiation factor 2 subunit 3 Y-linked (*Eif2s3y*) (Table S3). Other genes located in autosomes and upregulated in females were *Krt13,* Lymphocyte antigen 6 family member d (Ly6d), Small proline-rich protein 2a3 (Sprr2a3), Semaphorin 3a (Sema3a), Transmembrane serine protease 11 (Tmprss11) family and Catenin delta 2 (Ctnnd2). In the males some of the top upregulated were S100 calcium binding protein a4 (s100a4), Fk506 binding protein 5 (Fkbp5), Regulator of G-protein signaling-like protein (Rgcc), Carcinoembryonic antigen-related cell adhesion molecule 2 (Ceacam2) and Lymphocyte antigen 6 family member A (Ly6a).

Overall, the spatially resolved scRNAseq analysis reveals conserved epithelial subtypes in the mouse urethra and highlights sex-specific gene expression differences.

### Krt5^+^ cells are a self-renewing stochastic population maintaining homeostasis

To assess whether the molecular differences identified with the scRNAseq analysis translate into different cellular dynamics, we decided to trace the fate choices that accompany the homeostasis of the entire urethral epithelium. We first employ the *Krt5Cre-ERT2*;*Rosa26^YFP^* reporter mice (Figure 5A-B, S3A). Given that Krt5 is equally expressed in the basal and intermediate cells of the entire urethral epithelium of both sexes and that proliferative cells are present in greater proportion in the basal layer and with a minor presence in the intermediate layer (Figure S3B-C), the Krt5 promoter allowed us to study proliferative cell behavior and self-renewal capacities at the population level. Since all epithelial layers are Cd326^+^ and the basal cells express Cd49f in higher levels than the other cell layers (Figure S2B), these markers could be used in FACS to isolate the epithelial cells. To assess whether Krt5⁺ cells contribute to a self-sustaining compartment, we first monitored the evolution of YFP-labeled cells over a 24-week period (Figure 5C). We observed a 3-fold initial expansion in labeled cell fractions, followed by a clear plateau. From a theoretical perspective, this suggested that Krt5^+^ labels a representative basal population that self-maintains in a balanced manner, but expands over short time by giving rise to intermediate and luminal layers^33^ (Document S4). This also allowed us to estimate the proliferation rate of basal cells, by fitting the time-course data to a simple compartment model of basal, intermediate and luminal cells. From this fit, we extracted effective division rates of λ = 0.315/week or 22 days for females and λ = 0.209/week or 34 days for males (Figure 5C). These estimates reflect the average number of divisions per week per labeled cell contributing to the labeled pool. While recombination was initially higher in females, the long-term plateauing and fit-supported kinetics support the notion that Krt5⁺ cells behave as a self-renewing population with sex-specific division rates.

**Figure 5:**
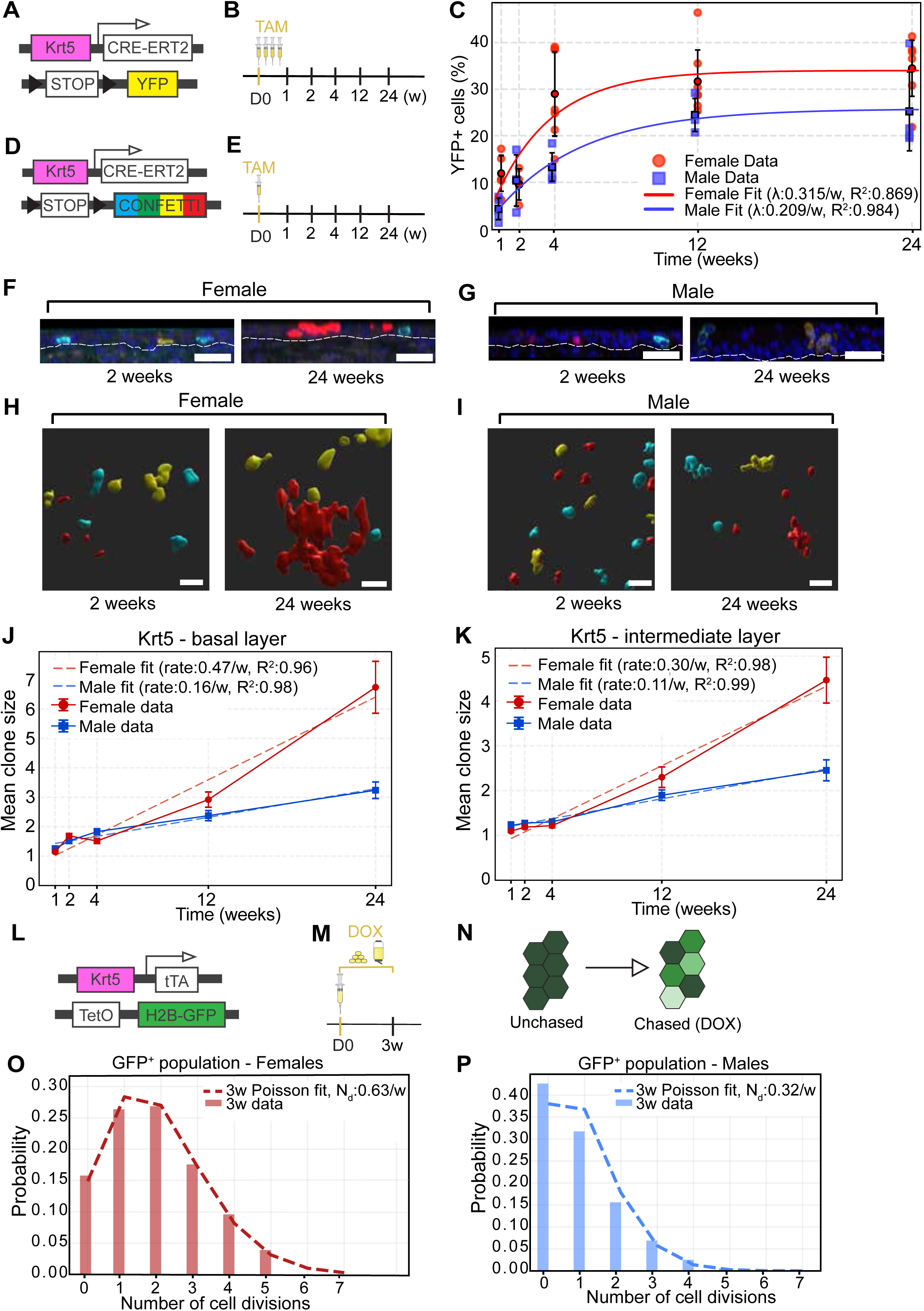
**Krt5 marks a self-sustained population**A. *Krt5Cre-ERT2;Rosa26^YFP^* mouse construct. B. Experimental strategy. *Krt5Cre-ERT2;Rosa26^YFP^* mice (n=29 females and 18 males) received four injections of tamoxifen in 4 consecutive days and the cells were chased for 1-24 weeks. C. Percentage of epithelial cells that are YFP^+^ in females (red) and males (blue) for the five timepoints. Data represented as average per mouse ± SD, each point representing flow cytometry data from one mouse. The mean of all mice of the same timepoint is highlighted with a black outline. D. *Krt5Cre-ERT2;Rosa26^Confetti^* mouse construct. E. Experimental strategy. *Krt5Cre-ERT2;Rosa26^Confetti^* mice (n=15 per sex) received an injection of tamoxifen at the start of the experiment and the cells were chased for 1-24 weeks. F. Orthogonal view of wholemount immunofluorescence staining of *Krt5Cre-ERT2;Rosa26^Confetti^*females for the 2 week and 24 weeks timepoints. Endogenous Confetti expression (CFP-cyan, YFP-yellow, RFP-red) and Hoechst (blue). Basement membranes are outlined with dashed lines. Scale bar: 30μm. G. Orthogonal view of wholemount immunofluorescence staining of *Krt5Cre-ERT2;Rosa26^Confetti^*males for the 2 week and 24 weeks timepoints. Endogenous Confetti expression (CFP-cyan, YFP-yellow, RFP-red) and Hoechst (blue). Basement membranes are outlined with dashed lines. Scale bar: 30μm. H. 3D cell volume visualization rendering of wholemount immunofluorescence staining on a female at 2 and 24 weeks. I. 3D cell volume visualization rendering of wholemount immunofluorescence staining on a male at 2 and 24 weeks. J. Quantification of *Krt5Cre-ERT2;Rosa26^Confetti^* data for the basal layer. Mean clone size for basal cells of females (red) and males (blue) are plotted against the 5 timepoints. The data have been fitted with a regression line and the coefficient of determination (R^2^) is highlighted. The slope of each fit is used to calculate the rate of division. K. Quantification of *Krt5Cre-ERT2;Rosa26^Confetti^* data for the intermediate layer. Mean clone size for intermediate cells of females (red) and males (blue) are plotted against the 5 timepoints. The data have been fitted with a regression line and the coefficient of determination (R^2^) is highlighted. The slope of each fit is used to calculate the rate of division. L. *Krt5tTA;tetO;H2B-GFP* mouse construct. M. Experimental strategy. *Krt5tTA;tetO;H2B-GFP* mice (n=7 females and 8 males) received an injection of doxycycline at the start of the experiment as well as doxycycline food and water for the whole three weeks chase period. N. Schematic of theoretical H2B-GFP color dilution in induced *Krt5tTA;tetO;H2B-GFP* mice. The color is diluted by 2 with each division of chased cells. After multiple cell divisions, a homogenous patch of cells in an unchased tissue (left) will take a heterogeneous appearance (right) in the chased (DOX) tissue. The color intensity will depend on how many times each cell divided (right). O. Poisson fit of female *Krt5tTA;tetO;H2B-GFP* mice experimental data for the 3w DOX chase. P. Poisson fit of male *Krt5tTA;tetO;H2B-GFP* mice experimental data for the 3w DOX chase.

Next, to provide insights into the fate behavior of individual cells, we turned to *Krt5Cre-ERT2*;*Rosa26^Confetti^* reporter mice and induced them with a very low dose of Tamoxifen to perform a clonal analysis, ensuring that we analyzed the progeny of individually labelled cells (Figure 5D-E, Table S5-S6). Even though the majority of the labelled cells started from the basal layer, the progeny of both female and male cells reached the luminal layer within the 24 weeks of tracing (Figure 5F-G). Following the labeling, the mean clone size of Krt5⁺ cells in the basal, intermediate and luminal layers was quantified over time in both sexes. First, we looked at the clone size distribution depending on proximal or distal region for both females and males and we found that the female regions had very similar rates while males were more diverse (Figure S3D-H). However, since comparing individual regions was challenging from a statistical perspective, and since we were interested in comparing the global scale dynamics between the different sexes, we pooled the regional data together. At the start of the tracing, the majority of the labelled cells were at the 1 cell clone size, but the clone size increased in both the basal and intermediate compartment as time progressed (Figure 5H-K), while clonal persistence decreased (Figure S3D). This is a hallmark of stochastic cell fate decisions leading to neutral drift, which predicts a linear increase in surviving clone size ⟨n(t)⟩ = 1 + λ_s_t/2, where λ_s_ is a symmetric division rate of basal cells. As this linear prediction could reproduce well the data in both males and females, we could estimate the symmetric division rates at λ_s_=0.47/week for females and λ_s_ =0.16/week for males. This confirmed the previous finding (Figure 5C) that female Krt5⁺ cells divide faster than their male counterparts. The difference in magnitude may reflect model assumptions (e.g., neutrality, compartment mixing) or technical distinctions between population- and clone-level measurements.

To confirm that the difference in the clone size between males and females was the result of a difference in the rate of cell proliferation, we performed pulse-chase experiments using the *Krt5tTA*;*tetO*;*H2B-GFP* reporter mouse^34^ (Figure 5L-M, S4A). In the absence of doxycycline (DOX), histone 2B domain–green fluorescent protein (H2B–GFP) fusion protein is expressed at a high level in a discrete peak of fluorescence in FACS (Figure 5N, S4A). Following DOX administration, the H2B–GFP signal decreases by a factor of two with each cell division (Figure 5M-N, S4A). Thus, the level of H2B-GFP label dilution represents the number of cell divisions accomplished by the basal Cd49f^+^ cells during the chase period, as quantified by FACS (Figure S4A). The H2B-GFP label dilution was used to estimate division numbers in Krt5⁺ basal cells at 3 weeks post-DOX administration for females and males (Figure 5O-P). We converted fluorescence intensities to inferred division counts and fit these distributions with a stochastic division model (Poisson distribution), where the mean N_d_ reflects the average number of divisions per cell. At 3 weeks, division distributions were well-fitted by a single, unimodal, Poisson distribution, with estimated rates of 0.63/week for females and 0.32/week for males (Figure 5O-P). These overall division rates are slightly higher than the symmetric division rates extracted from the clonal analysis, consistent with a number of other studies showing stochastic decisions between symmetric divisions, asymmetric divisions and symmetric differentiation^35–39^. Together, these data suggest that progeny of Krt5^+^ cells populate all epithelial cell layers and that female Krt5^+^ cells proliferate around twice as fast as male Krt5^+^ cells.

### Krt14^+^ cells behave with a faster yet limited self-renewal potential

Given that Krt14 is only expressed in the basal layer for both females and males and that in other epithelia such as the skin epidermis and the bladder urothelium is considered a marker of stem cells, we decided to trace Krt14^+^ cells, a subpopulation of the Krt5^+^ cells. To study the Krt14 progeny, we turned to the *Krt14-rtTA;TetOCRE;Rosa26^mTmG^*mouse model and first assessed the labelling of the Krt14-derived cells at the population level overtime (Figure 6A-D, S4B). The percentage of labelled basal cells remained constant for 24 weeks after the initial labelling, indicating that the Krt14^+^ population is self-renewing. However, when looking at the shape of the clones and even though the same number of cells were recombined between sexes, the clone morphology appears different (Figure 6C). We noticed that the mGFP^+^ clones were mostly single cells and more dispersed in the females, while in the males, the clones were more packed (Figure 6C). Given that clonal shape and dispersion has been recently suggested to be related to the physical state of the tissue – with tissue fluidity favoring cellular re-arrangements and clonal dispersion^40–46^, we reasoned that female tissues might be more fluid. However, we did not find a difference between the shapes of individual female and male cells that could explain this behavior (Figure S4C-D).

**Figure 6:**
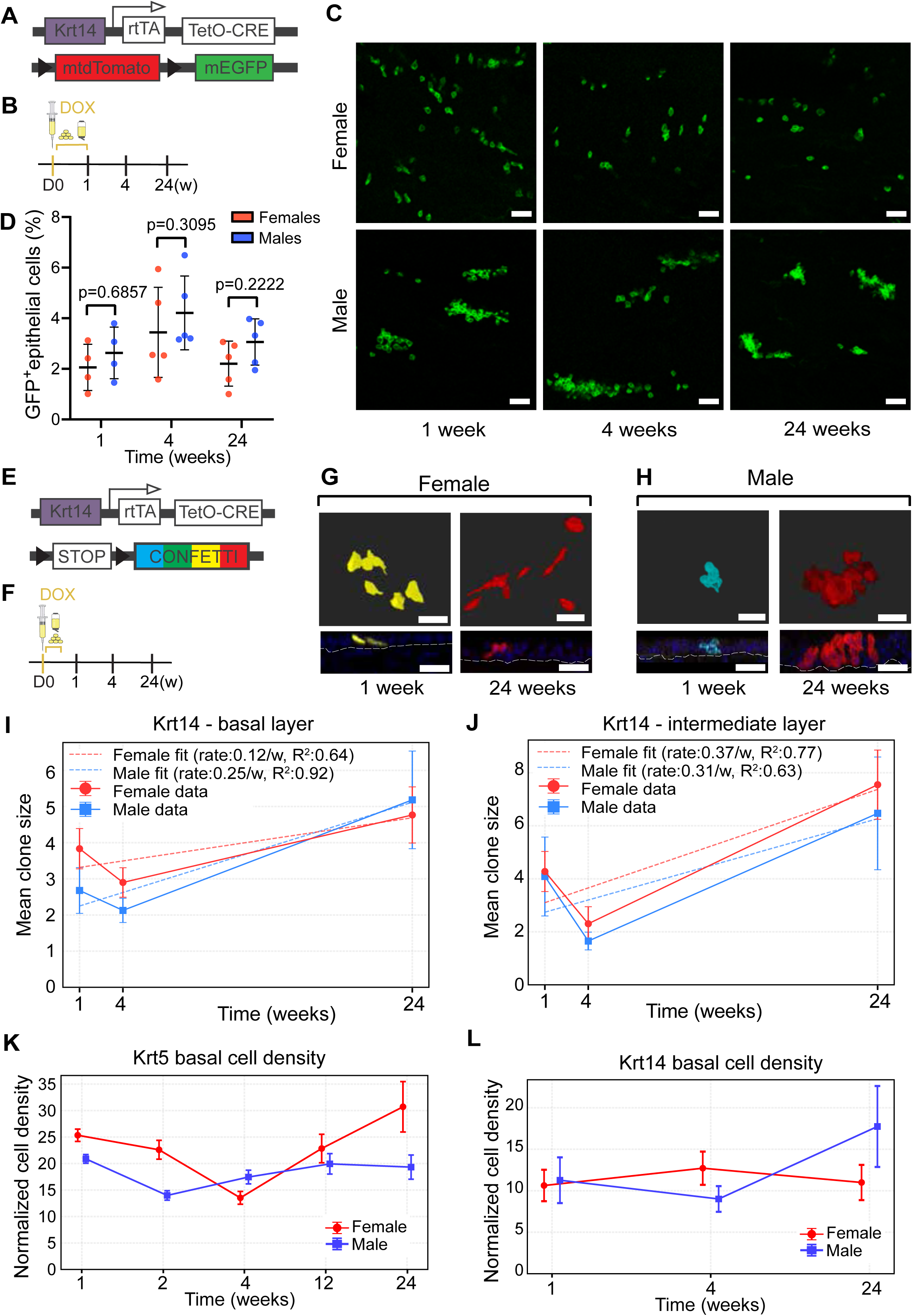
**Krt14 marks a self-sustained population**A. *Krt14-rtTA;TetOCRE;Rosa26^mTmG^* mouse construct. B. Experimental strategy. *Krt14-rtTA;TetOCRE;Rosa26^mTmG^* mice (n=14 per sex) received an injection of doxycycline at the start of the experiment as well as doxycycline food and water for 1 week. The chase period lasted 1-24 weeks. C. Maximum intensity projection of wholemount immunofluorescence staining for females and males at 1, 4 and 24 weeks. Endogenous mGFP expression shown in green. Scale bar: 50μm. D. Percentage of epithelial cells that are GFP+ in females (red) and males (blue) at 1, 4 and 24 weeks. Data represented as mean ± SD, each point representing flow cytometry data from one *Krt14-rtTA;TetOCRE;Rosa26^mTmG^*mouse. Statistical test is Mann-Whitney U test. E. *Krt14-rtTA;TetOCRE;Rosa26^Confetti^* mouse construct. F. Experimental strategy. *Krt14-rtTA;TetOCRE;Rosa26^Confetti^* mice (n=9 per sex) received an injection of doxycycline at the start of the experiment as well as doxycycline food and water for 4 days. The chase period lasted 1-24 weeks. G. 3D cell volume visualization rendering of wholemount immunofluorescence staining of a female at 1 and 24 week. Below the visualizations, the corresponding orthogonal views are displayed with the basement membranes marked with dashed yellow line. Endogenous Confetti expression (CFP-cyan, YFP-yellow, RFP-red) and Hoechst (blue). Scale bar: 30μm. H. 3D cell volume visualization rendering of wholemount immunofluorescence staining of a male at 1 and 24 week. Below the visualizations, the corresponding orthogonal views are displayed with the basement membranes marked with yellow dashed line. Endogenous Confetti expression (CFP-cyan, YFP-yellow, RFP-red) and Hoechst (blue). Scale bar: 30μm. I. Quantification of *Krt14-rtTA;TetOCRE;Rosa26^Confetti^* data for the basal layer. Mean clone size for basal cells of females (red) and males (blue) are plotted against the 3 timepoints. The data have been fitted with a regression line and the coefficient of determination (R^2^) is highlighted. The slope of each fit is used to calculate the rate of division. J. Quantification of *Krt14-rtTA;TetOCRE;Rosa26^Confetti^* data for the intermediate layer. Mean clone size for basal cells of females (red) and males (blue) are plotted against the 3 timepoints. The data have been fitted with a regression line and the coefficient of determination (R^2^) is highlighted. The slope of each fit is used to calculate the rate of division. K. Normalized basal cell clone density (cells/area) of female (red) and male (blue) *Krt5Cre-ERT2;Rosa26^Confetti^* for all the timepoints. Data represented as mean ± SD. L. Normalized basal cell clone density (cells/area) of female (red) and male (blue) *Krt14-rtTA;TetOCRE;Rosa26^Confetti^* for all the timepoints. Data represented as mean ± SD.

Thus, to understand how these differences occur, we traced the dynamics of single cells by performing a quantitative clonal analysis, taking advantage of the *Krt14-rtTA;TetOCRE;Rosa26^Confetti^* mice model (Figure 6E-H, S5A, Table S5-S6). Interestingly, the cells show the same behavior as seen in the *Krt14-rtTA;TetOCRE;Rosa26^mTmG^* model with the males exhibiting more clustered clones with greater distance between them while the female clones are more elongated and dispersed (Figure 6G-H, S5A). Strikingly, there was a marked difference in the short-term clonal output when compared to Krt5^+^ cells. Although we know that with this model we always start with labelled cells in the basal compartment, we see a large increase in the intermediate compartment, and we can already see luminal cells at the 1w timepoint for both females and males (Figure 6G-J, S5B). This data is in contrast with the *Krt5Cre-ERT2*;*Rosa26^Confetti^*reporter mice where the first female luminal clones appear at the 4w timepoint (the male ones appear earlier probably due to lower tissue stratification). Overall, at 1w, Krt14^+^ clones already consist of an average of 4-5 cells (compared to only 1-2 in Krt5^+^), suggesting a considerably high proliferation rate (Figure S5C-D). This was not due to clonal mixing of multiple labelled clones in the same location as there was a considerable distance between clones of a given color (Figure S5A). Even more surprisingly, when tracing Krt14^+^ clones over time, we found no evidence of clonal loss or clone size increase, again in stark contrast with the neutral drift dynamics observed for Krt5^+^ (Figure S5B-D). We found the same trend as in the Krt5-tracing where the proximal region has a higher proliferation rate (Figure S5E-H). When comparing the clone density for Krt5- and Krt14-labelled progeny, the Krt5 data show an initial decrease in density for both females and males, which then reaches a plateau after the 4w timepoint (Figure S5F) while the Krt14 data for both females and males follow a plateau for the whole 24 weeks (Figure S5B). When looking at the labelled cell density, the data follows a plateau for both the Krt5 and Krt14 until the end of the experiment at the 24w timepoint (Figure 6K-L). When looking at the cumulative probability for the Krt5^+^ and Krt14^+^ clones, the higher proliferative capacity of the Krt14^+^ clones within the same timeframe is evident (Figure S6A-D). The most parsimonious explanation for the Krt14^+^ behavior is the classical proliferative unit traditionally proposed long-time ago in skin, where asymmetrically dividing stem cells produce fast-dividing and short-lived progenitors – explaining how clones both reach a large clone size early and then plateau^47,48^.

## Discussion

Here we provide a thorough description of the urethral epithelium and characterize regional and sex differences between epithelial cells. Moreover, we generated the first scRNAseq dataset of the full-length adult mouse urethra for both females and males, which can be used to understand the cellular heterogeneity and complexity of this tissue.

Firstly, when comparing the proximal to the distal urethral epithelium, we observe large changes in the shape of the urethral epithelium and stratification in both sexes. Secondly, we provide evidence for differences in the urethral epithelium between sexes, something that has not been described before in literature. We find that the females have a more stratified epithelium and a higher division rate than the males in all the tracing experiments we conducted. On average, female cells proliferate faster than male cells for both the Krt14^+^ and Krt5^+^ cells. We also identified sex differences in the expression of the epithelial markers such as Krt13 and Krt20.

Comparing to the other epithelia such as the skin, intestinal and vaginal epithelium, the urethral epithelium exhibits a significantly lower turnover rate^15,49–51^. However, the urethral epithelium has a higher turnover of around 22 days in females and 34 days in males, compared to the bladder epithelium which has an average turnover of 200 days under homeostasis^52,53^. Previous results have been reported for the homeostatic conditions in the adult mouse bladder: Papafotiou et al., have shown that the Krt14^+^ cells, a small population of basal Krt5^+^ cells, can generate all three (basal, intermediate and luminal) cell layers^54^, while Wang et al. and Gandhi et al., reported that Krt5^+^ cells have not been able to generate luminal cells during a 2 weeks tracing^55,56^. However, a 2 weeks tracing is a very short timeframe in comparison to the homeostatic bladder turnover rates of 200 days. In our tracing of the equivalent cells in the urethras of *Krt5Cre-ERT2*;*Rosa26^Confetti^*mice, we detected basal-derived luminal cells in stratified epithelium at 4 weeks in the females and at 2 weeks in the males. Our findings indicate that both Krt5^+^ and Krt14^+^ basal cells can generate all cell layers in the urethra of both sexes under homeostatic conditions. Interestingly, the Krt14^+^ cells were able to achieve this much faster than the Krt5^+^ progeny.

To interpret the dynamic behaviors of urethral progenitors beyond static imaging or marker expression, we implemented mathematical models that quantitatively capture population turnover and clonal growth across sexes and basal cell types. By fitting various independent datasets within a shared modeling strategy, we identified consistent sex differences in cell division rates and uncover distinct renewal strategies for Krt5⁺ and Krt14⁺ cells. Indeed, Krt5⁺cells showed behavior consistent with neutral competition, where clones expand or are lost over time through stochastic symmetric divisions. This is characteristic of an equipotent, self-renewing progenitor population, with fate choices balanced at the population level. In contrast, Krt14⁺ clones expanded rapidly at early time points and then stabilized in both size and density. This behavior does not fit the predictions of neutral drift and instead suggests that Krt14^+^ cells behave as long-lived and asymmetrically dividing progenitors, which constantly produce a short-lived, transit-amplifying population. It is tempting to speculate that this might correspond to a division of labor, with a slow population ensuring long-term homeostasis while a fast population enables fast responses to physiological perturbations experienced by the urethral epithelium. This is reminiscent of findings in the skin epithelial where upon wound healing, a specific population of asymmetrically dividing cell produces fast progenies^57^. These findings point to meaningful functional heterogeneity within the basal layer and highlight how clonal modeling can uncover renewal strategies that cannot be inferred from marker expression alone.

Looking into the future, there is clear potential to expand the clonal modeling framework to capture more of the biological variability we observe. While the current models distinguish effectively between neutral and hierarchical behaviors, they assume that all cells within a given lineage behave in the same way, which could be relaxed to consider heterogeneity or fate conversion. Models that account for fluctuations in fate choice or division timing may also help us better understand how clones persist or are lost. With more densely sampled time courses and longer-term lineage tracing, the modeling framework could also be used to investigate clonal dominance, stability, and potential age-related changes in progenitor output.

Altogether, the modeling approaches used here support and deepen our experimental findings. They show that the urethra is maintained by multiple progenitor types with distinct behaviors, and that these strategies vary between sexes. The reason behind these epithelial sex differences is currently unknown and outside the scope of this manuscript. Our work establishes the urethra as a powerful model system in its own right - one that is accessible, well-structured, and well-suited for studying fundamental questions in epithelial homeostasis and stem cell biology.

## Supporting information

Table S1: Counts from the scRNAseq data

Table S2: Marker genes of epithelial subclusters from scRNAseq data

Table S3: Differentially expressed genes between sexes in epithelial cells from scRNAseq data

Document S4: Supplementary note on cell behaviour mathematical modelling

Table S5: Quantification of female and male Krt5Cre-ERT2;Rosa26Confetti and Krt14-rtTA;TetOCRE;Rosa26Confetti

Table S6: Number of clones per normalized area of female and male Krt5Cre-ERT2;Rosa26Confetti and Krt14-rtTA;TetOCRE;Rosa26Confetti

## Acknowledgements

We thank Mia Kristine Grønning Høg and Ida Marie Egeland Larsen from BRIC Histology and Microscopy Core Facility for help with sample preparation, Javier Martin Gonzalez and the Core Facility for Transgenic Mice at Copenhagen University for mice transfer, rederivation and support and the staff of the animal facilities at Copenhagen University. We also thank Gelo Dela Cruz from the reNEW Flow Cytometry Platform, Jutta Bulkescher, Jonas Viswalingam Bagge and Antonios Georgantzoglou from the reNEW Microscopy Facility, as well as Heike Wollmann, Adrija Kalvisa and Magali Michaut from the reNEW Genomics Platform for technical expertise, support, and use of instruments. We sincerely thank all other Aragona lab members for their feedback and engaging discussions.

This study has been supported by the Novo Nordisk Foundation Project Grant (NNF20OC0064992). The Novo Nordisk Foundation Center for Stem Cell Medicine (reNEW) is supported by the Novo Nordisk Foundation (NNF21CC0073729).

## Author contributions

I.K. and M.A. designed the experiments and performed data analysis; I.K. performed most of the experiments; O.B.A. helped with experiments and performed data analysis; U.R.G. and E.H. performed cell behavior modelling analysis; D.A. performed single-cell RNA sequencing bioinformatic analysis; M.A. supervised the work and acquired funding; I.K., U.R.G., O.B.A., E.H. and M.A. wrote the manuscript with suggestions from D.A.; All authors have proofread and approved the final version of the manuscript.

## Data availability

Single-cell RNA sequencing raw data are publicly available (GEO: GSE299898).

---

**Supplementary Figure 1:**
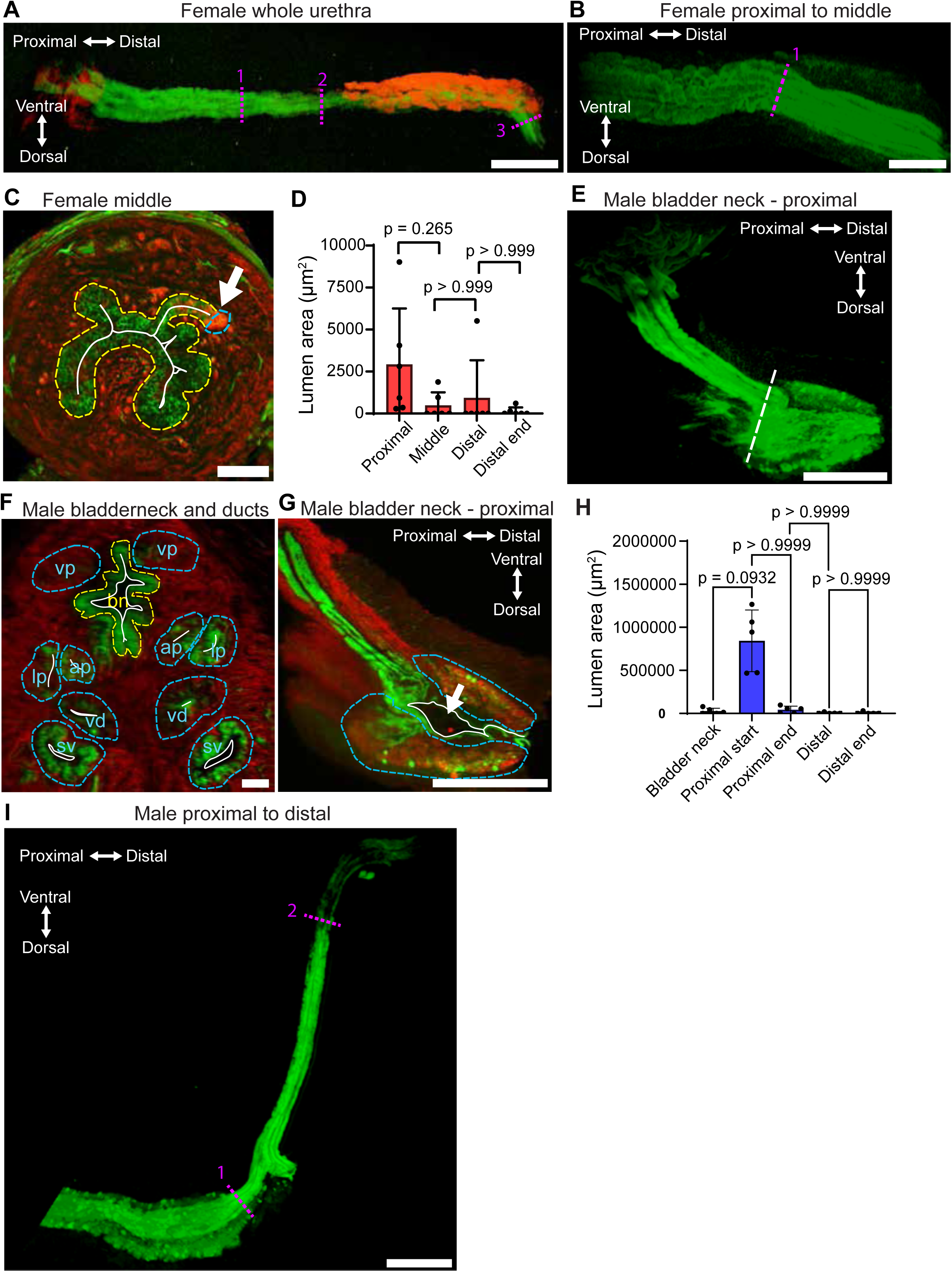
3D visualization of female and male urethra where tissue transitions are evident. A. 3D visualization of a *Krt5Cre-ERT2*;*Rosa26^mTmG^* female urethra, chased for 26 weeks. Proximal and distal directions are indicated in white. Scale bar: 1500μm. B. Close-up of a *Krt5Cre-ERT2*;*Rosa26^mTmG^* female proximal to middle region at the first tissue transition, chased for 26 weeks. Proximal and distal directions indicated in white. Scale bar: 500μm. C. Cross-section of a *Krt5Cre-ERT2*;*Rosa26^mTmG^* female urethra, chased for 26 weeks, at the start of the distal region. The white arrow indicates a location where a duct of the Skene’s glands is merging with a urethral fold. Scale bar: 100μm. D. Quantification of the female urethra lumen at the proximal, middle, distal and distal end regions, n=6 mice. Data represented as mean ± SD. Statistical test is Friedman test with Dunn’s multiple comparisons tests. E. 3D visualization of the bladder neck connecting to the start of the proximal urethra in a male *Krt5Cre-ERT2*;*Rosa26^mTmG^* chased for 40 weeks. Proximal and distal directions indicated in white. White dashed line indicates cross section shown in Figure S1F. Scale bar: 1500μm. F. Cross-section of a male *Krt5Cre-ERT2*;*Rosa26^mTmG^* urethra chased for 40 weeks at the first tissue transition. The urethra (ur) is running in parallel to the ducts connecting the ventral prostate (vp), anterior prostate (ap), lateral prostate (dp), vas deferens (vd) and seminal vesicles (sv). Scale bar: 100μm. G. Longitudinal section of E, with the large proximal lumen indicated with a white arrow. Proximal and distal directions indicated in white. Scale bar: 1500μm. H. Quantification of male urethral lumen at the start and end of the bladder neck, proximal and distal urethra, n=5 mice. Data represented as mean ± SD. Statistical test is Friedman test with Dunn’s multiple comparisons tests. I. 3D visualization of remaining proximal, membranous and distal urethra in a *Krt5Cre-ERT2*;*Rosa26^mTmG^* male chased for 40 weeks. Proximal and distal directions indicated in white. Scale bar: 1500μm. A-C, E-G, I. Recombined *Krt5Cre-ERT2*;*Rosa26^mTmG^*urethras. Membrane GFP in green, membrane tdTomato in red. Ventral direction is the top of each image while dorsal direction is the bottom of each image in this figure. Basement membranes are marked in yellow and glands marked in blue. Tissue transitions are marked magenta.

**Supplementary Figure 2:**
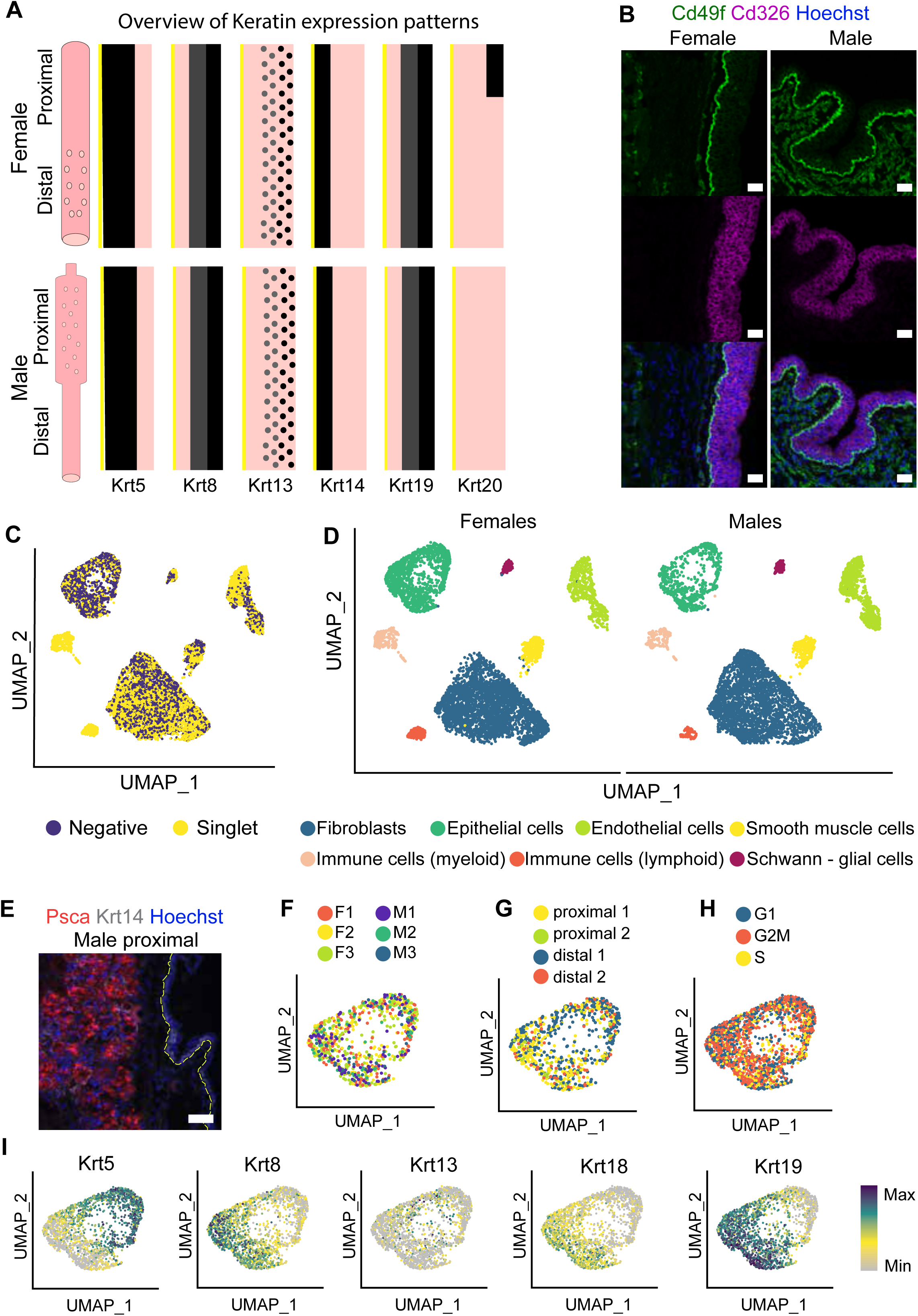
Strategy for the single-cell RNA sequencing experiment and distribution of the cells within the clusters. A. Overview schematic of keratin expression (Krt5, Krt8, Krt13, Krt14,, Krt19 and Krt20) in female and male urethras depending on the region (proximal or distal) and cell layer (basal, intermediate, luminal). A uniform expression is indicated with a solid line while a salt-and-pepper expression is indicated with a dotted line. Strong expression is indicated with a dark black line; weaker expression is indicated with a gray line and absent expression is indicated with a pink line. The basement membrane is marked with a yellow line. B. Immunofluorescence staining on a longitudinal section for Cd49f (green), Cd326 (magenta) and nuclei (blue) in representative female and male urethras. Scale bar: 30μm. C. Distribution of hashed cells (singlet in yellow) and non-hashed cells (negative in blue) in the UMAP of all cells, sexes combined. D. UMAP of the distribution of the seven different cell clusters based on sex. Fibroblasts are in blue, epithelial cells in dark green, endothelial cells in light green, smooth muscle cells in yellow, immune cells of myeloid lineage in beige, immune cells of lymphoid lineage in orange and Schwann-glial cells in magenta. E. Immunofluorescence staining on a longitudinal section for the gland marker Psca (red), Krt14 (gray) and nuclei (Hoechst). Basement membrane marked with yellow dashed line. Scale bar: 60μm. F. UMAP plot colored by the origin sample distribution form the six animals (females F1-F3 and males M1-M3) used in the experiment within the epithelial cluster. G. UMAP plot colored by the origin sample distribution form the four regions (proximal 1 and 2, distal 1 and 2) used in the experiment within the epithelial cluster. H. Predicted cell-cycle phases within the epithelial cluster, assigned using Seurat’s CellCycleScoring function and visualized in the UMAP. Cells in G1 are in blue, cells in G2/M are in orange and cells in S phase are in yellow. I. UMAP plot colored by normalized gene expression values for the indicated keratin gene and in the epithelial cluster. Gene expression is visualized as a color gradient going from gray to violet with gray as indicator of no expression and violet as indicator of maximal expression.

**Supplementary Figure 3:**
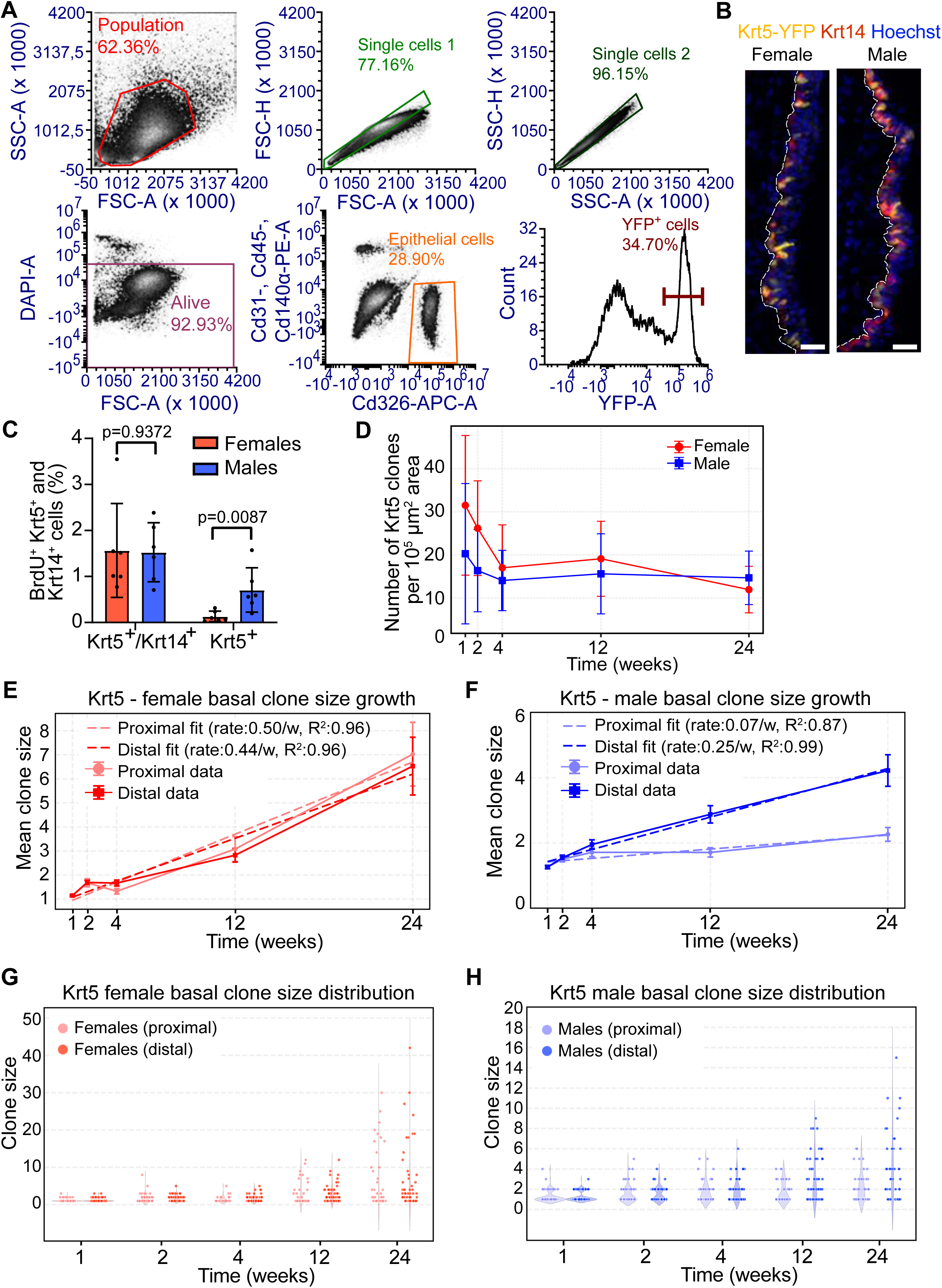
Flow cytometry can be used to segregate Krt5 basal cells which are shown to be proliferative. A. Flow cytometry strategy for isolating live YFP+ epithelial cells (cells that are YFP^+^, Cd326^+^, Cd31^-^, Cd45^-^, Cd140a^-^) for quantitative analysis. B. Immunofluorescence staining on longitudinal sections of female and male *Krt5Cre-ERT2;Rosa26^YFP^*mice for endogenous Krt5-YFP (yellow), Krt14 (red) and nuclei (Hoechst). Basement membranes are marked with a white dashed line. Scale bar: 30μm. C. Percentage of BrdU^+^ cells in female (red) and male (blue) urethras after 4 hours of chase, n=6 mice per sex. Data is shown as mean fraction of BrdU^+^ cells ± SD that were either Krt5^+^ or Krt14^+^Krt5^+^. There were no BrdU^+^ cells that were Krt14^+^ Krt5^-^. Each point is showing the average counts per image for each mouse. Statistical test is Mann-Whitney U test. D. *Krt5Cre-ERT2;Rosa26^Confetti^* clone survival over the 24 weeks for females (red) and males (blue). Data represented as mean ± SD. E. Quantification of *Krt5Cre-ERT2;Rosa26^Confetti^* female basal layer data for proximal (light red) and distal (dark red) region. The mean clone size for basal cells is plotted against the 5 timepoints. The data have been fitted with a regression line and the coefficient of determination (R^2^) is highlighted. The slope of each fit is used to calculate the rate of division. F. Quantification of *Krt5Cre-ERT2;Rosa26^Confetti^* male basal layer data for proximal (light blue) and distal (dark blue) region. The mean clone size for basal cells is plotted against the 5 timepoints. The data have been fitted with a regression line and the coefficient of determination (R^2^) is highlighted. The slope of each fit is used to calculate the rate of division. G. Basal clone size distribution of *Krt5Cre-ERT2;Rosa26^Confetti^*females based on proximal (light red) or distal (dark red) region for all 5 timepoints. H. Basal clone size distribution of *Krt5Cre-ERT2;Rosa26^Confetti^* males based on proximal (light blue) or distal (dark blue) region for all 5 timepoints.

**Supplementary Figure 4:**
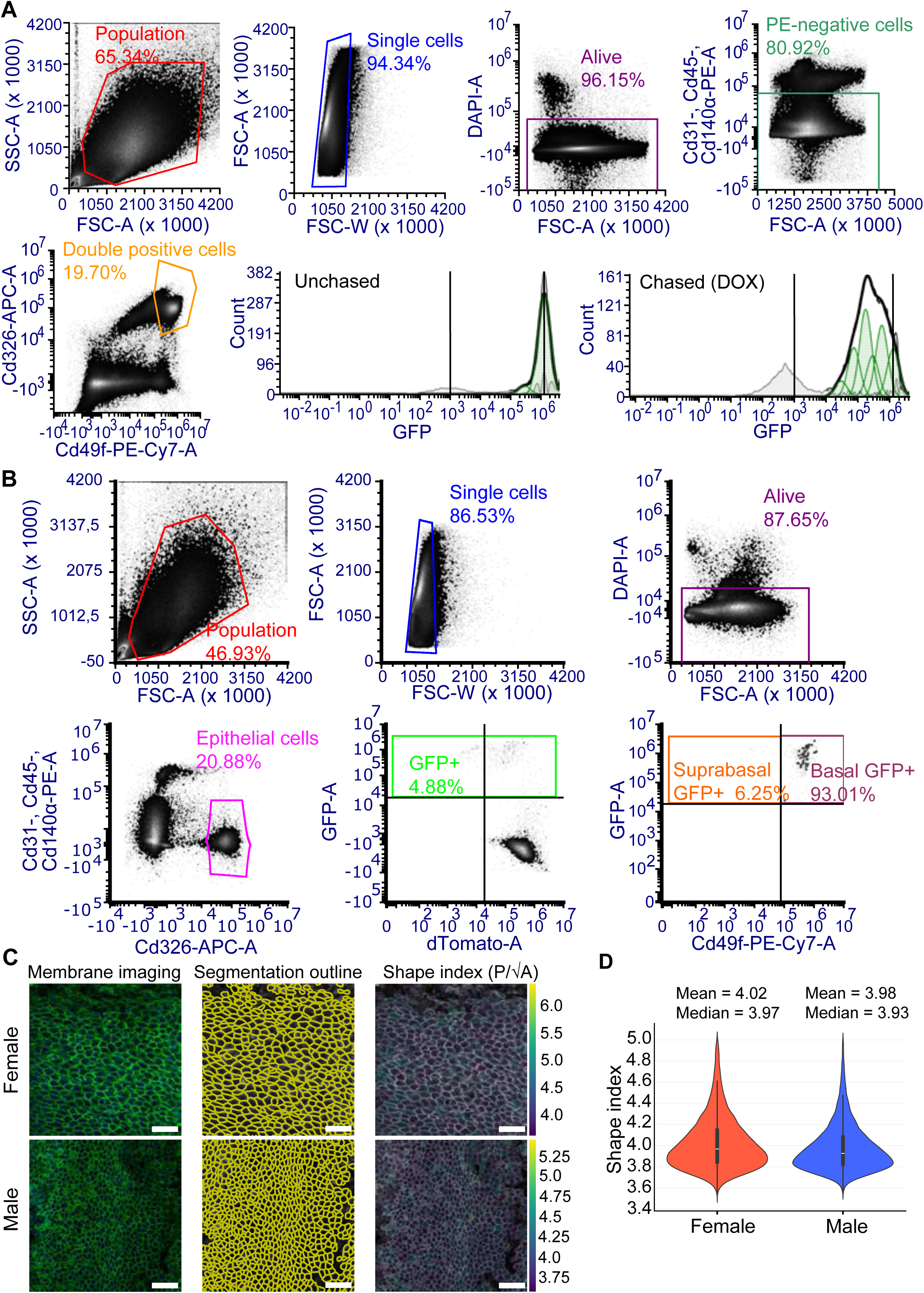
Flow cytometry and cell shape strategies for analyzing the behavior of Krt5 and Krt14 cells. A. Flow cytometry strategy for isolating live basal epithelial cells (cells that are Cd49f^+^, Cd326^+^, Cd31^-^, Cd45^-^, Cd140a^-^) from *Krt5tTA;tetO;H2B-GFP* mice. These cells are then allocated based on their GFP expression into cell divisions for quantitative analysis. The 3w-chased cells show multiple peaks compared to the unchased control. The multiple peaks and vertical lines highlight the strategy for allocating GFP^+^ cells (GFP-Area signal: 10^3^-10^6^) into cell divisions. B. Flow cytometry strategy for isolating live GFP^+^ epithelial cells (the GFP^High^, Cd326^+^, Cd31^-^, Cd45^-^, Cd140a^-^cells) from *Krt14-rtTA;TetOCRE;Rosa26^mTmG^* for quantitative analysis. Basal and suprabasal GFP^+^ cells were separated using Cd49f. C. Representative images from female (top) and male (bottom) *Krt5Cre-ERT2*;*Rosa26^mTmG^*epithelium used for shape quantification. Left: membrane imaging using a combination of cell membrane (mGFP and mtdTomato in green) and nuclear (Hoechst in blue) markers. Middle: segmentation outlines generated using both membrane and nuclear channels to improve boundary detection. Right: shape index maps, computed as p/√A (perimeter over square root of area), with individual cells color-coded by index value. Scale bar: 50 µm. D. Distribution of cell shape indices across sexes. Both male and female tissues show overlapping distributions, with only a minor difference in mean shape index (female mean = 4.02; male mean = 3.98). Despite slight numerical variation, this difference is small and unlikely to reflect meaningful mechanical or topological differences.

**Supplementary Figure 5:**
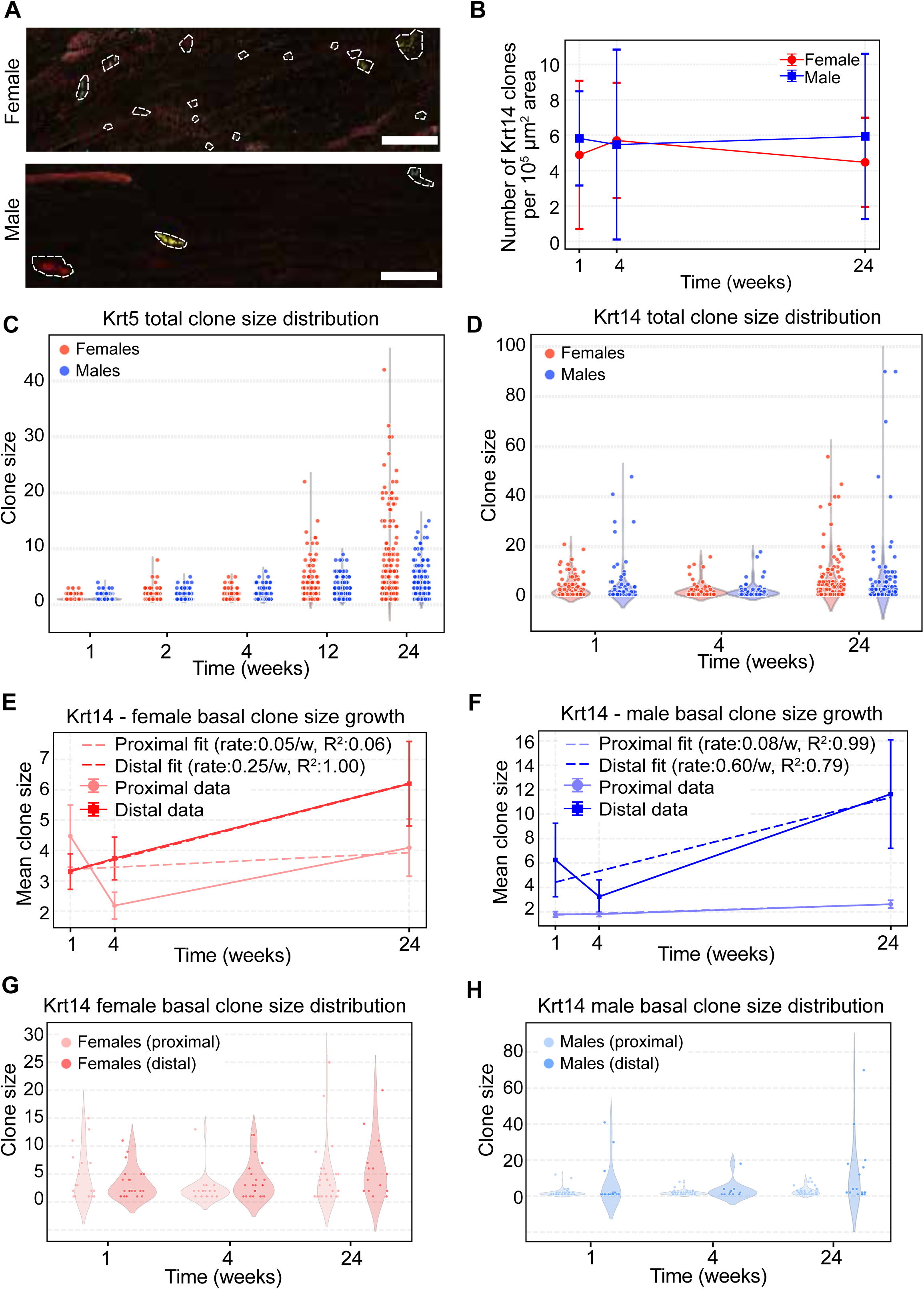
Krt14 basal cells are proliferative and behave differently to Krt5 cells. A. Wholemount immunofluorescence staining on a representative female and male *Krt14-rtTA;TetOCRE;Rosa26^Confetti^*urethra at the 1w timepoint. The clones visible in the images are marked with dashed lines. The average distance between each clone to its closest neighboring clone is 172μm for the female and 1185μm for the male. Scale bar: 400μm. B. *Krt14-rtTA;TetOCRE;Rosa26^Confetti^* clone survival over the 24 weeks for females (red) and males (blue). Data represented as mean ± SD. C. Total clone size distribution of *Krt5Cre-ERT2;Rosa26^Confetti^* females (red) and males (blue) for all 5 timepoints. D. Total clone size distribution of *Krt14-rtTA;TetOCRE;Rosa26^Confetti^* females (red) and males (blue) for all 3 timepoints. E. Quantification of *Krt14-rtTA;TetOCRE;Rosa26^Confetti^* female basal layer data for proximal (light red) and distal (dark red) region. The mean clone size for basal cells is plotted against the 3 timepoints. The data have been fitted with a regression line and the coefficient of determination (R^2^) is highlighted. The slope of each fit is used to calculate the rate of division. F. Quantification of *Krt14-rtTA;TetOCRE;Rosa26^Confetti^* male basal layer data for proximal (light blue) and distal (dark blue) region. The mean clone size for basal cells is plotted against the 3 timepoints. The data have been fitted with a regression line and the coefficient of determination (R^2^) is highlighted. The slope of each fit is used to calculate the rate of division. G. Basal clone size distribution of *Krt14-rtTA;TetOCRE;Rosa26^Confetti^*females based on proximal (light red) or distal (dark red) region for all 3 timepoints. H. Basal clone size distribution of *Krt14-rtTA;TetOCRE;Rosa26^Confetti^*males based on proximal (light blue) or distal (dark blue) region for all 3 timepoints.

**Supplementary Figure 6:**
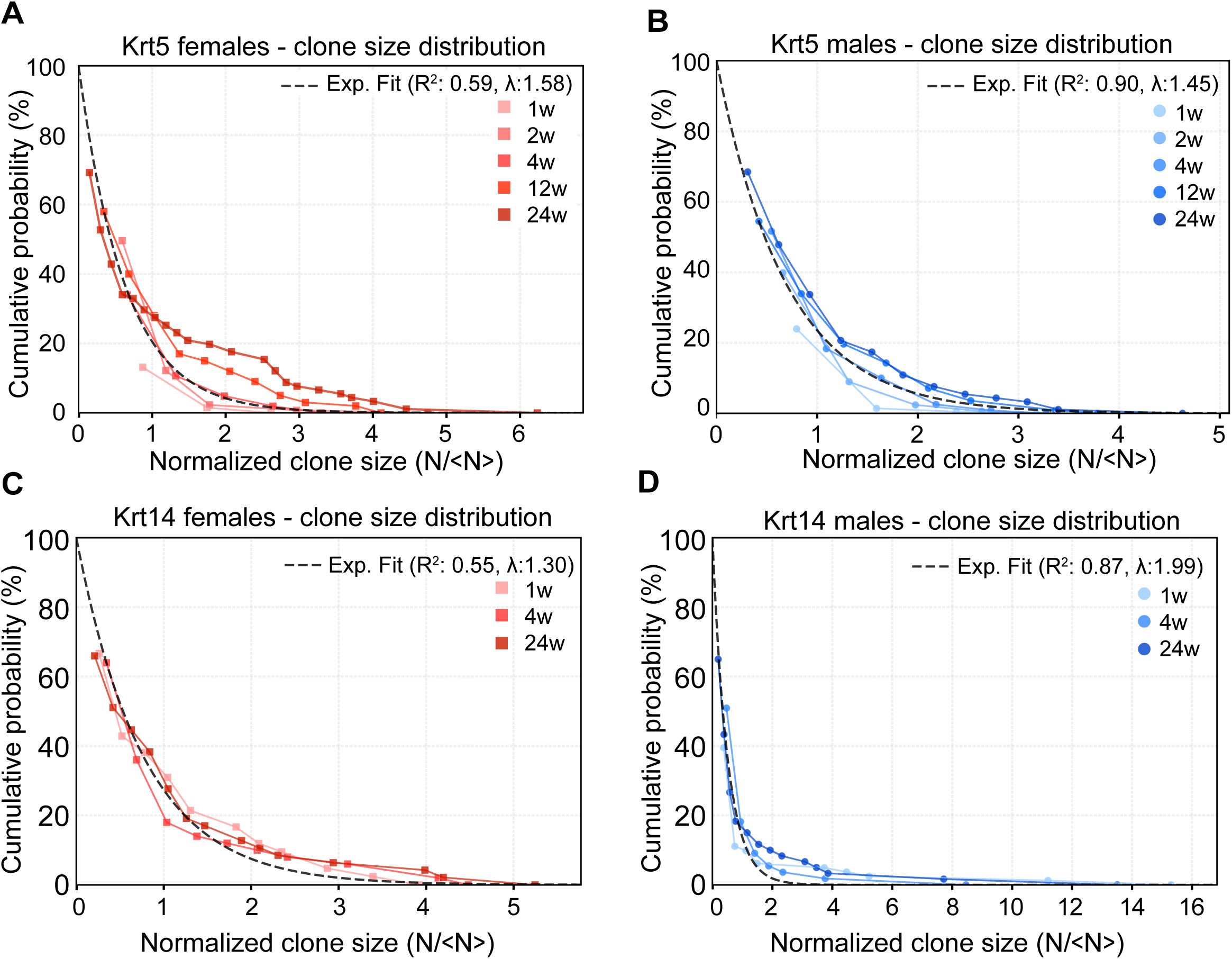
**The Krt14 and Krt5 progeny have different behaviors**A. Cumulative probability of female *Krt5Cre-ERT2;Rosa26^Confetti^* clones for all the timepoints. B. Cumulative probability of male *Krt5Cre-ERT2;Rosa26^Confetti^* clones for all the timepoints. C. Cumulative probability of female *Krt14-rtTA;TetOCRE;Rosa26^Confetti^*clones for all the timepoints. D. Cumulative probability of male *Krt14-rtTA;TetOCRE;Rosa26^Confetti^* clones for all the timepoints.

## Materials & Methods

### Mice

All animal experiments were approved by the Danish Animal Experimental Inspectorate (Dyreforsøgstilsynet) under license number 2021-15-0201-00861 and conducted in accordance with Directive 2010/63/EU of the European Parliament and the Council on the protection of animals used for scientific purposes. The breeding and experimental activities on mice were conducted at the Faculty of Health and Medical Sciences of Copenhagen University, at the Department of Experimental Medicine.

The mice were maintained under a 12-hour light-dark cycle with lights turning on at 6AM. Temperature was kept constant at 22°C and the animals had ad libitum access to food and water. All mice are of mixed background and group housed (unless separation was deemed necessary). The adult mice were sacrificed by cervical dislocation, while pups by decapitation. Mice used in this study were composed of males and females with mixed genetic background and aged from 60 to 320 days. The exact age of the mice is specified in each experiment. No statistical methods were used to predetermine sample size and the experiments were not randomized. The investigators were not blinded to allocation during experiments and outcome assessment. All the transgenic mice that were used [FVB.Cg-Tg(Krt5-tTA), Tg(tetO-H2b-GFP)1JH, FVB.Cg-Tg(KRT5-cre/ERT2)2Ipc/JeldJ, B6.129X1-Gt(ROSA)26Sor^tm1(EYFP)Cos^/J, B6.129-*Gt(ROSA)26Sor^tm^*^4^*^(ACTB−tdTomato,−EGFP)Luo^*/J), B6.129-*Gt(ROSA)26Sor^tm^*^1^*^(CAG-^*^Brainbow.1)Cle^/J, Tg(KRT14-rtTA)F42Efu and B6.Cg-Tg(tetO-cre)1Jaw/J] were described previously^34,58–64^.

### Induction and lineage tracing of *Krt5Cre-ERT2;Rosa26^mTmG^*

P60-P90 male and female *Krt5Cre-ERT2;Rosa26^mTmG^* mice were injected intraperitoneally with Tamoxifen powder (Sigma-Aldrich #T5648) dissolved in a solution of 10% ethanol and 90% corn oil (Thermo Scientific Chemicals #405435000) to create a 25mg/ml Tamoxifen solution. A dose of 5mg Tamoxifen per 30g mouse weight (or 200μl of the tamoxifen solution per 30g mouse weight), once per day for three consecutive days was used. Mice were then left for 24-40 weeks until sacrifice to allow sufficient turnover of the epithelium.

### Induction and lineage tracing of *Krt5Cre-ERT2;Rosa26^YFP^*

P60-P90 male and female *Krt5Cre-ERT2;Rosa26^YFP^* mice were injected intraperitoneally Tamoxifen powder (Sigma-Aldrich #T5648) dissolved in a solution of 10% ethanol and 90% corn oil (Thermo Scientific Chemicals #405435000) to create a 25mg/ml Tamoxifen solution. A dose of 5mg Tamoxifen per 30g mouse weight (or 200μl of the tamoxifen solution per 30g mouse weight), once per day for four consecutive days was used. Mice were then sacrificed at 1 week-, 2 week-, 4 week-, 12 week- and 24 week-timepoints.

### Induction and lineage tracing of *Krt5Cre-ERT2;Rosa26^Confetti^*

P60-P90 male and female *Krt5Cre-ERT2;Rosa26^Confetti^*mice were injected intraperitoneally with Tamoxifen powder (Sigma-Aldrich #T5648) dissolved in a solution of 10% ethanol and 90% corn oil (Thermo Scientific Chemicals #405435000) to create a 10mg/ml Tamoxifen solution. A single dose of 2mg Tamoxifen per 30g mouse weight (or 200μl of the tamoxifen solution per 30g mouse weight) was used. Mice were then sacrificed at 1 week-, 2 week-, 4 week-, 12 week- and 24 week-timepoints.

### Induction and lineage tracing of *Krt14-rtTA;TetOCRE;Rosa26^mTmG^*

P60-P90 male and female *Krt14-rtTA;TetOCRE;Rosa26^mTmG^*mice were injected intraperitoneally at D0 with 1,2mg/30g mouse weight doxycycline hyclate (Sigma-Aldrich #D9891) dissolved in saline (Fresenius Kabi #B306443/01) and treated with doxycycline-infused food (Bio-Serv #S3888) and water for 7 days. The doxycycline-infused water was prepared by dissolving 0.1% w/v doxycycline hyclate (Sigma-Aldrich #D9891) in drinking water and mixed until completely dissolved. Food and water were changed every 2-3 days. The mice were sacrificed at 1 week-, 4 week- and 24 week-timepoints.

### Induction and lineage tracing of *Krt14-rtTA;TetOCRE;Rosa26^Confetti^*

P60-P90 male and female *Krt14-rtTA;TetOCRE;Rosa26^Confetti^*mice were injected intraperitoneally at D0 with 1,2mg/30g mouse weight doxycycline hyclate (Sigma-Aldrich #D9891) dissolved in saline (Fresenius Kabi #B306443/01) and treated with doxycycline-infused food (Bio-Serv #S3888) and water for 4 days. The doxycycline-infused water was prepared by dissolving 0.1% w/v doxycycline hyclate (Sigma-Aldrich #D9891) in drinking water and mixed until completely dissolved. Food and water were changed every 2-3 days. The mice were sacrificed at 1 week-, 4 week- and 24 week-timepoints.

### BrdU proliferation experiments

P60-P90 male and female SWISS (Janvier Labs) mice were injected once with a dose of 1,5mg/30g mouse weight of 5-Bromo-2′-deoxyuridine (BrdU) (Sigma-Aldrich #B5002) dissolved in filtered Phosphate Buffered Saline (PBS). The mice were dissected 4 hours after the injection.

### H2B-GFP proliferation experiments

P60-P90 male and female *Krt5tTA;tetO;H2B-GFP* mice were injected intraperitoneally at D0 with 1,2mg/30g mouse weight doxycycline hyclate (Sigma-Aldrich #D9891) dissolved in saline (Fresenius Kabi #B306443/01) and treated with doxycycline-infused food (Bio-Serv #S3888) and water for the whole duration of the experiment. The doxycycline-infused water was prepared by dissolving 0.1% w/v doxycycline hyclate (Sigma-Aldrich #D9891) in drinking water and mixed until completely dissolved. Food and water were changed every 2-3 days. Control (unchased) mice received a saline injection instead of doxycycline injection at D0 and had ad libitum access to normal food and water. The mice were sacrificed at the 3 week timepoint.

### Tissue preparation for histological analysis

Paraffin samples were prepared by fixing the tissues in 4% w/v Paraformaldehyde (PFA) in milliQ water (Thermo Fisher Scientific #28908) overnight at 4°C, washed in 70% ethanol and stored in 70% ethanol until they were dehydrated through a graded series of ethanol and embedded in paraffin. The organs were cut into 4μm-thick sections using a HM 355S Automatic Microtome (Thermo Fisher). The sections were de-paraffinized, then rehydrated through ethanol series and lastly stained Hematoxylin and Eosin (H&E).

### Whole-mount and section preparation for immunostaining

For whole-mount tissue preparation, the dissected urethras were opened with a fine scissor and fixed while flattened for 2 hours at RT in 4% w/v PFA prior to washing 3 times in PBS. For whole-mount staining, the tissues were blocked for 5 hours at RT with the whole-mount blocking buffer [1% w/v Bovine Serum Albumin (BSA) (Sigma-Aldrich #A9418), 5% v/v horse serum (Gibco #26050088), 0.8% v/v Triton X-100 (Sigma-Aldrich #X100) in PBS]. Right after, the tissues were incubated with primary antibodies for two days at 4°C. The third day, tissues were washed 3 times in PBST [PBS with 0.2% v/v TWEEN® 20 (Sigma-Aldrich #P1379)] for 20 minutes, followed by incubation with secondary antibodies for 2 hours at RT. Then, slides were washed 2 times in PBST and one time in PBS before mounting with 2.5% w/v 1,4-Diazabicyclo[2.2.2]octane (Sigma-Aldrich #D27802) in Glycerol Mounting Medium (Dako #C0563). For the BrdU staining, post-fixation tissues were first incubated for 30 minutes in pre-heated 1N hydrochloric acid at 37°C and washed 3 times for 10 minutes in filtered PBS before whole-mount staining described as above began.

For generating slides for immunofluorescence, the dissected urethras were placed in Tissue-Tek O.C.T. Compound (Sakura #4583), frozen on dry ice pellets and cut into 8μm thick sections using the CM1950 cryostat (Leica Biosystems). For slide staining, the tissue sections were fixed for 10 minutes at RT in 4% w/v PFA, followed by 3 PBS washes for 5 minutes. Then, sections were blocked for 1 hour at RT with the slide blocking buffer [1% w/v BSA (Sigma-Aldrich #A9418), 5% v/v horse serum (Gibco #26050088), 0.2% v/v Triton X-100 (Sigma-Aldrich #X100) in PBS]. For antibody staining, the slides were incubated with primary antibodies overnight at 4°C and the next day, after being washed 3 times in PBS for 5 minutes, they were incubated with secondary antibodies for 1 hour at RT. Slides were washed again 3 times in PBS before mounting with 2.5% w/v 1,4-Diazabicyclo[2.2.2]octane (Sigma-Aldrich #D27802) in Glycerol Mounting Medium (Dako #C0563).

The following primary antibodies were used: anti-Krt5 (Biolegend #905901), anti-Krt8 (DHSB Hybridoma Bank #TROMA-I), anti-Krt13 (Abcam #ab92551), anti-Krt14 (Invitrogen #PA5-16722), anti-Krt14 (Origene #BP5009), anti-Krt19 (DHSB Hybridoma Bank #TROMA-III), anti-Krt20 (My BioSource #MBS833195), anti-Krt20 (Progen #GP-K20), anti-BrdU (Abcam #6326), anti-Psca (Invitrogen #PA5-115799), anti-Lgals7 (Abcam #ab10482), anti-Nipal1 (My BioSource #MBS9205311), anti-Sprr1a (Invitrogen #PA5-110423). The following secondary antibodies (Jackson Immunoresearch) were used: anti-rabbit, anti-chicken, anti-guinea pig and anti-rat Rhodamine Red-X, Alexa Fluor 488, Alexa Fluor A647 and Cy5. Nuclei were stained with Hoechst (Invitrogen #H3570).

### Lightsheet tissue preparation and microscopy

After sufficient turnover of the epithelium, *Krt5Cre-ERT2;Rosa26^mTmG^*mice were sacrificed and their urethras were dissected with bladder attached. They were subsequently fixed for 5 hours in 4% w/v PFA at RT. Bladders were removed after 2 hours of fixation. Samples were then washed 3 times in filtered PBS for 20 minutes and left in decoloration medium [25% w/v quadrol (Sigma-Aldrich #102-60-3) in milliQ water] at 37°C shaking overnight. Samples were then washed 3 times in filtered PBS for 20 minutes at 37°C shaking, and subsequently incubated in 50% v/v tetrahydrofuran (Sigma-Aldrich #5895700100) in milliQ water for 20 hours shaking at RT. Samples were then washed for 4 hours in milliQ water, changing the water every hour. Samples were then incubated in whole-mount blocking buffer with Sytox Deep Red nuclear stain (1:2000, Invitrogen #S11380) shaking at RT overnight. The clearing medium EZ View was prepared according to a previously described method^65^. Finally, samples were cleared in EZ View for 24 hours, before being embedded in 2% w/v low melting point agarose (Thermo Fisher #16500100) in milliQ water, before re-incubation in EZ View medium for an additional 24 hours. EZ View medium was refreshed every 12 hours to ensure a stable refractive index. Samples were imaged on a Lightsheet 7 (ZEISS). Deconvolution, stitching and chromatic aberration correction was done using Huygens Professional (Scientific Volume Imaging). Image analysis was performed in Imaris version 10 (Oxford Instruments) and Fiji (ImageJ).

### Flow cytometry

Urethras were dissected, minced using scissors and placed in digestion solution to incubate at 37°C for 2 hours on a shaker. The digestion solution is 7,5 ml Hanks’ Balanced Salt solution (HBSS) (Thermo Fisher Scientific #14025100) supplied with 0.25% w/v Collagenase I (Sigma-Aldrich C0130), 2.5mM Calcium chloride (Thermo Scientific Chemicals #423525000) and 0.005% w/v DNase I (Roche #11284932001). The digestion was stopped by the addition of 75μl of 0.5M Ethylenediaminetetraacetic acid (EDTA) (Thermo Fisher Scientific #15575020), the suspensions were filtered through 70μm strainers, then 40μm strainers and then centrifuged to obtain cell pellets. The final pellets were re-suspended in chelated FACS (cFACS) buffer [PBS-Ca-Mg supplied with 2% v/v chelated Fetal Bovine Serum (FBS)]. Chelated FBS was prepared by dissolving 20% w/v Chelex 100 chelating Resin (Biorad #1422842) in FBS (Gibco # 10082147).

To distinguish different cell types during flow cytometry analysis, cells were stained for 30 minutes with conjugated antibodies, followed by washing with cFACS buffer. The antibodies used were CD45-PE (1:1200, Invitrogen #12-0451-82), CD31-PE (1:1200, BD Biosciences #553373), CD49f-PE-Cy7 (1:2000, Invitrogen #25-0495-82), EPCAM-APC (1:1200, Thermo Fisher #17-5791-82) and PDGFRα-PE (1:200, Invitrogen #12-1401-81). Cells were also stained with 1.7μM 4’,6-diamidino-2-phenylindole (DAPI) (Invitrogen #D1306) and filtered through 40μm strainers right before analysis which was performed using Cytek Aurora cytometer (Cytek Biosciences).

All the flow cytometry data were analyzed with SpectroFlo (Cytek Bio) and FCS Express (De Novo software). To allocate GFP^+^ cells of *Krt5tTA;tetO;H2B-GFP* mice to a specific number of cell divisions, the data were mapped with the Proliferation Fit from FCS Express for up to 5 divisions.

### Single-cell RNA sequencing library preparation

SWISS mice (Janvier Labs) were used and urethras were cut in 4 equal length pieces for males and 2 equal pieces for females and digested separately. Cell pellets were prepared as described above. Instead of incubation with fluorescent antibodies, each sample was incubated with hashing antibodies (1:100, BioLegend TotalSeq™-A03XX anti-mouse Hashtag) for 20 minutes in order to be able to be distinguished when pooled for sequencing. Each sample was also stained with 0.3mM DAPI. Sorting was performed using the FACSymphony S6 sorter (BD Biosciences) with a 70μm nozzle. Live cells were sorted using DAPI into the bottom of Eppendorf tubes, same sex samples pooled together during sorting.

The 2 samples (female and male) were immediately processed for single-cell sequencing and libraries were generated using the Chromium Next GEM Single Cell 3’ Kit v3.1 (10X Genomics) as per manufacturer’s instructions. Hashtag barcode libraries, we prepared as described before^66,67^. The material and oligonucleotide sequences for generating the hashtag barcode libraries can be found in the Key Resources Table. All libraries were diluted to 4 nM and the female and male cDNA libraries were pooled with their respective hashtag libraries at a ratio of 95:5. Then, the female and male total libraries were pooled at a 1:1 ratio to be sequenced together using a P2 100 kit. Sequencing was done on the NextSeq2000 sequencer (Illumina).

### Single-cell RNA sequencing data analysis

Raw sequencing data generated with the Illumina NextSeq 2000 platform were demultiplexed using *bcl-convert* (v4.1.7-2) and aligned to the mouse genome (mm10, GENCODE vM23/Ensembl 98) or HTO tag sequences using *Cell Ranger* (v6.0.1). On average, 25 million reads per sample were sequenced. The *Cell Ranger --multi* workflow, with the *--include-introns* option, was used to align and quantify RNA and HTO reads for downstream analysis. Raw UMI count data were analyzed in R (v4.0.3) using the *DropletUtils* and *Seurat* packages. Droplets with abnormally high RNA or HTO counts were excluded. For the female sample, only A0301–A0306 hashtags were retained. The *barcodeRanks* and *emptyDrops* functions from *DropletUtils* were applied to RNA and HTO count data to distinguish empty droplets from true cells. Only droplets identified as non-empty in either dataset were retained. Final cell counts were 7,214 for the male sample and 8,216 for the female sample, matching expected recovery rates.

Low-quality cells were excluded using the *isOutlier* function, applying thresholds of three MADs below and five MADs above the mean, with maximum limits of 15,000 UMIs and 6,000 detected genes. This resulted in the removal of 785 female and 910 male cells. Sample assignment was performed using three complementary methods: *hashedDrops*, *HTODemux*, and *MULTIseqDemux*. Results were consolidated using a majority-rule framework to identify high-confidence singlets, exclude doublets, and flag discordant assignments. Demultiplexing accuracy was assessed using a t-SNE plot of CLR-normalized HTO data.

Damaged cells with elevated mitochondrial or ribosomal gene expression were excluded using the *adjOutlyingness* function. Ambient RNA contamination was estimated with *decontX* function, and cells with contamination levels above 25% were removed. Cells with high hemoglobin gene expression and Y-chromosome transcripts in female samples were also excluded. After filtering, 79% of female and 76% of male cells were retained, primarily removing damaged and contaminated cells.

Next, built-in S-phase and G2/M-phase marker genes were mapped to mouse (*Mus musculus*) orthologs using the *gProfiler2*, and cell cycle phases were assigned with Seurat’s *CellCycleScoring* function.

Filtered count matrices were merged, retaining genes expressed in at least 10% of cells from the smallest sample. Highly variable genes (top 25%) were identified using the *FindVariableFeatures* function. UMI counts were normalized with *SCTransform*, regressing out mitochondrial gene expression. Principal component analysis (PCA) was performed using *RunPCA*. The optimal number of PCs was determined based on *ElbowPlots*, dimensional heatmaps, and the *findCutoff* algorithm, selecting between 10 and 15 PCs depending on dataset characteristics. SCT-normalized data were subsequently integrated using the Harmony algorithm via *Seurat::IntegrateLayers*.

Cell clustering was optimized to balance granularity and biological interpretability. An SNN graph was constructed from Harmony-integrated dataset using *FindNeighbors* with cosine distance, followed by Louvain clustering. Uniform Manifold Approximation and Projection (UMAP) was applied to facilitate low-dimensional visualization of high-dimensional transcriptomic data. The *k.neigh* parameter was set to twice the expected number of cell types. Cluster-specific marker genes were identified using the *FindMarkers* function with the Poisson test. To ensure a balanced number of cells per cell type, the dataset was downsampled to the smallest number of cells per cell type per sample. Marker genes were cross-referenced with cell-type enriched gene lists from the Human Protein Atlas Single Cell resources https://www.proteinatlas.org/humanproteome/single+cell/single+cell+type. Only upregulated genes meeting the following criteria were retained: Adjusted p-value < 0.001; Log2 fold-change > 2.5; Expression in ≥75% of cells within the cluster. Top 10 genes based on log2 Fold change were plotted in the Heatmap. Cell identities were assigned after evaluating marker genes in the context of scientific literature.

Differential gene expression analysis between distal and proximal regions, as well as male and female samples, was performed using a pseudo-bulk DESeq2 approach and the *FindMarkers* function. The p-value threshold was set at the 5th percentile of the p-value distribution, with an adjusted p-value cutoff of 0.05. The fold-change threshold was defined as ±1.5 standard deviations, with a minimum absolute fold change of 0.5. Only genes expressed in at least 25% of cells in a cell type in one of the samples were retained.

### Imaging and analysis

Images of wholemounts and some cryosections were acquired using the confocal microscopes LSM 780 (ZEISS), Cell Observer (ZEISS), SP8 (Leica Biosystems) and Stellaris (Leica Biosystems). Images of the rest of the cryosections were acquired with the widefield microscope DM5500 (Leica Biosystems). Images of paraffin sections were acquired with the NanoZoomer slidescanner (Hamamatsu Photonics).

Images of paraffin sections were manually quantified for muscle thickness, epithelium thickness and number of cell layers using the NDP.view2 (Hamamatsu Photonics) software. Images of *Krt5Cre-ERT2;Rosa26^Confetti^* and *Krt14-rtTA;TetOCRE;Rosa26^Confetti^* mice were manually quantified and annotated using the softwares ZEN (ZEISS) and LASX (Leica Biosystems). For these, the images were quantified for number of clones per image, color per clone, number of cells per clone and location of each cell per clone in reference to the three epithelial cell layers. The number of clones per image were converted to number of clones per standard area (=10^5^μm^2^) to normalize the different frame size of the 3 different confocal microscopes that were used over the years. At least 30 clones per region (proximal, distal, gland epithelium) per mouse were quantified and the location of each image was annotated on the tile scan of each mouse.

Orthogonal views of z-stacks for wholemount staining were created using the Fiji software. Images of BrdU experiments were quantified using the software ZEN (ZEISS), LAS X (Leica Biosystems) and Fiji with the Cell Counter plugin. For these, maximum intensity projections were quantified for the number of Krt5^+^ cells, Krt14^+^ cells, Krt5^+^/ Krt14^-^cells, Krt5^-^/ Krt14^+^ cells and any co-localization with BrdU^+^. At least 3 orthogonal z-stacks per region per mouse were imaged and quantified, and the location of each image was annotated on the tile scan of each mouse. The male proximal and distal urethras were divided into 5 regions each while the female was divided into 6 regions.

The Krt14 coverage was quantified by producing tile scans of Krt5-, and Krt14-stained wholemounts on the LSM780 microscope using a combination of tiling and z stack generation. The Krt13 and Krt20 coverage was quantified by producing tile scans of Krt5-, Krt13- and Krt20-stained wholemounts using the navigator implementation on the SP8 microscope where the epithelium was brought into focus on every tile. Using Fiji, background intensity was thresholded, and binary images were created of Krt13, Krt14, Krt20 and Krt5. For the Krt14 images, the setting ‘‘Enhance Local Contrast (CLAHE)’’ was used in Fiji to equalize the intensities over the samples to avoid elimination of low intensity (or slightly out of focus) cells during thresholding. Using Krt5 to outline the entire epithelium, pixels were quantified and Krt13-, Krt14- and Krt20-coverages were normalized relative to the Krt5 coverage.

### Imaris rendering

3D volume visualizations of *Krt5Cre-ERT2;Rosa26^Confetti^*and *Krt14-rtTA;TetOCRE;Rosa26^Confetti^* mice were performed with the Imaris software (Oxford Instruments) by using the surface rendering tool to mask the single colors of the confetti reporter and produce 3D volumes.

### Cell shape analysis

Fluorescent microscopy images of mouse urethral epithelium were acquired as multi-channel.lif files and processed using a custom image analysis pipeline written in Python. Cell segmentation was performed using the CellPose 3 algorithm^68^, following a de-noising pre-processing step to enhance membrane signal clarity. A dual-channel strategy was employed, combining nuclear staining (Hoechst, channel 0) with mGFP and mtdTomato membrane markers (maximum intensity projection of channels 2 and 3) to improve boundary detection. For each segmented cell, morphological parameters were extracted, including cell area, perimeter, and shape index, calculated as perimeter divided by the square root of the area (perimeter/√area). Physical measurements were calibrated using the microscope’s pixel size metadata, with area reported in μm² and perimeter in μm. Images with fewer than ten identifiable cells were excluded from the analysis. Cells were grouped by experimental condition (sex and urethral region), and outliers with shape index values greater than 5 were excluded. Summary statistics, including mean and median values, were calculated for each group. Morphological data distributions were visualized using violin plots with embedded box plots to convey both variation and central tendencies. All image processing and analysis were conducted using a Python-based pipeline.

### Quantification and statistical analysis

Normality was checked when deciding for parametric tests. For experiments where n=3, we assumed that the data did not follow a normal distribution, so the appropriate nonparametric test was used instead. The exact p values are indicated above each comparison on each graph, differences between groups with p<0.05 were statistically significant. Figure legends indicate sample sizes, statistical tests and relevant multiple comparison corrections used. Graphs and statistical tests were generated using GraphPad Prism 10 and Python.

### Declaration of generative AI and AI-assisted technologies in the writing process

During the preparation of this work the authors used Copilot in order to correct grammar and shorten the summary. After using this tool, the authors reviewed and edited the content as needed and take full responsibility for the content of the publication.

### Supplemental files information

Table S1: Counts from the single-cell RNA sequencing data.

Table S2: Marker genes of epithelial subclusters from single-cell RNA sequencing data.

Table S3: Differentially expressed genes between sexes in epithelial cells from single-cell RNA sequencing data.

Document S4: Supplementary note on cell behavior mathematical modelling.

Table S5: Quantification of female and male *Krt5Cre-ERT2;Rosa26^Confetti^*and *Krt14-rtTA;TetOCRE;Rosa26^Confetti^*.

Table S6: Number of clones per normalized area of female and male *Krt5Cre-ERT2;Rosa26^Confetti^* and *Krt14-rtTA;TetOCRE;Rosa26^Confetti^*.

## Supplementary Note

### Quantitative analysis of progenitor dynamics

To systematically dissect the renewal strategies of urethral progenitor populations, we implemented a quantitative framework integrating multiple complementary analyses. Specifically, we examined: (i) clonal dynamics to distinguish neutral versus hierarchical fate behavior, (ii) sex-specific differences in population-level replacement kinetics, and (iii) direct division-rate measurements using H2B-GFP label dilution.

### Krt5^+^ population-level dynamics

To interpret the time evolution of YFP^+^ cell fractions in *Krt5Cre-ERT2;Rosa26^YFP^* mice (Figure 5C), we modeled epithelial maintenance using a simplified three-compartment framework. In this model, Krt5^+^ basal progenitors divide and give rise to intermediate and luminal cells, which are progressively replaced over time. Let *B*(*t*), *I*(*t*), and *L*(*t*) denote the number of labeled Basal, Intermediate, and Luminal cells, respectively. Their dynamics are governed by:

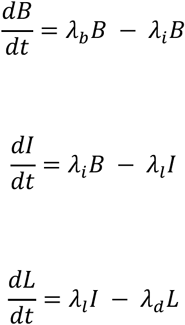

Here, *λ_b_* denotes the *total division rate* of basal cells, *λ_i_* is the *basal-to-intermediate differentiation rate*, *λ_l_* represents the *intermediate-to-luminal differentiation rate*, while *λ_d_* accounts for the *loss of luminal cells*, for instance due to cell shedding. Similar compartmental modeling framework, grounded in neutral drift theory, was previously applied by Doupé et al. to describe progenitor dynamics in the esophageal epithelium ^1^. While our study includes clonal lineage tracing, this model specifically describes *population-level replacement* inferred from cumulative YFP labeling. For instance, this whole compartment dynamics is not sensitive to whether basal divisions are symmetric or asymmetric, as it represents the global fluxes between compartments.

To analyze tissue-wide labeling kinetics, we focused on the total fraction of YFP^+^ epithelial cells over time:

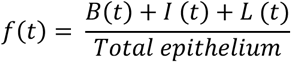

Under homeostatic conditions, the number of cells in each compartment remains constant, implying balanced fluxes. In particular, homeostasis requires that the rate at which basal cells divide equals the rate at which they differentiate into intermediate cells, i.e., *λ_b_* = *λ_i_*. To further simplify the system, and based on histological observations that all three compartments contain comparable numbers of cells at steady state, we assumed that:

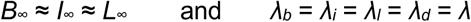

This approximation does not imply neutral drift — which concerns symmetric fate behavior at the clonal level ^2^ — but reflects a regime of uniform, balanced flux between compartments at the population level. Under these assumptions, the cumulative labeling dynamics simplify to a first-order saturation process with nonzero initial condition:

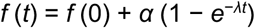

where *f* (0) is the initial labeled fraction of basal cells, and *α* captures the eventual contribution of these cells to the suprabasal layers. Since only basal cells are labeled at induction, *f* (0) represents the baseline label and the exponential term reflects upward flux into intermediate and luminal compartments. In this framework, the long-term labeling fraction reaches:

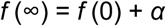

If compartments are of similar size and uniformly supplied by basal progenitors, we expect *α* ≈ 2*f* (0), yielding a final labeled fraction roughly threefold higher than the initial value.

Fitting this model to the experimental data yielded a good match across all timepoints. Female urethras exhibited a faster effective turnover rate (*λ* ≈ 0.3 week^−1^), while males showed a slower rate (*λ* ≈ 0.2 week^−1^). The fitted values of *f* (0) and *α* were both slightly higher in females, consistent with a combination of greater initial recombination efficiency and more efficient label propagation into suprabasal compartments.

Together, these results suggest that labeled Krt5^+^ progenitors progressively populate the epithelium via steady upward transitions, and that sex-specific differences in effective replacement rates underlie the observed labeling kinetics. While the model abstracts away clonal variability and assumes homogeneous behavior, it provides a useful coarse-grained view of epithelial turnover before accounting for cell-to-cell heterogeneity.

### Krt5^+^ division rate estimation from H2B-GFP label dilution

To independently quantify division rates of Krt5^+^ basal progenitors, we analyzed histone H2B-GFP label dilution at 3 weeks post-labeling in *Krt5tTA;tetO;H2B-GFP* mice (Figure 5L–P, S4A). In this system, H2B-GFP is expressed constitutively in basal cells in the absence of doxycycline (DOX), resulting in a strong, uniform nuclear signal. Upon DOX administration, new H2B-GFP production ceases, and the fluorescence intensity in each cell decreases by half with every cell division. Therefore, the extent of GFP signal dilution after a fixed chase period reflects the number of divisions each cell has undergone.

Fluorescence intensities measured by flow cytometry were converted to inferred division counts, assuming ideal halving of the H2B-GFP signal per mitosis. We then constructed histograms of cell division counts and fit these with a Poisson distribution:

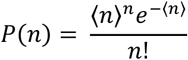

where *n* is the number of cell divisions, and ⟨*n*⟩ is the mean number of divisions per cell over the chase period. This model assumes that divisions are stochastic and independent, with a constant probability per unit time — conditions under which division events follow a Poisson process. Such a model is commonly used to describe population-level heterogeneity in proliferation under homeostatic conditions^3^.

The fitted Poisson distributions captured the general shape and spread of the experimental data (Figure 5O–P), consistent with a single, stochastic mode of proliferation in the Krt5^+^ population. From the fitted values of ⟨*n*⟩, we estimated the mean weekly division rates by dividing by the 3-week chase duration. Female basal cells exhibited an average division rate of approximately 0.63 week^−1^, while male cells showed a slower rate of 0.32 week^−1^.

While the Poisson model provides a minimal description of division count variability, it does not capture the full complexity of the data — particularly in females, where the distribution appears broader and more skewed. These differences suggest potential heterogeneity in proliferation kinetics that may not be fully described by a single rate parameter. Moreover, since the H2B-GFP readout reflects only the number of divisions and not the fate of daughter cells, this analysis does not distinguish between symmetric self-renewal, asymmetric division, or differentiation outcomes. Rather, it quantifies net proliferative activity within the basal Krt5^+^ pool.

Together, these findings confirm that Krt5^+^ progenitors are proliferative and that female cells divide more frequently than male counterparts, confirming the observed sex-specific differences in clonal expansion and epithelial turnover.

### Krt5^+^ clone size dynamics under neutral drift

To investigate cell fate behavior at the single-cell level, we performed long-term clonal lineage tracing in *Krt5Cre-ERT2;Rosa26^Confetti^* mice (Figure 5D–E). A low dose of tamoxifen was used to induce sparse recombination, allowing us to follow the expansion of individual Krt5^+^-derived clones over time. Although most labeled cells were initially located in the basal compartment, their progeny progressively reached intermediate and luminal layers in both sexes over the 24-week tracing period (Figure 5F–G).

Clone size was quantified separately in basal, intermediate, and luminal compartments at multiple timepoints, for both females and males. Initial regional analyses showed that female clone sizes were relatively uniform between proximal and distal urethral regions, whereas male clone sizes exhibited greater heterogeneity (Figure S3E–H). While these observations suggest potential spatial differences in clonal behavior—particularly in males—the limited number of clones captured per region constrained our ability to perform robust statistical comparisons across regions and timepoints. Moreover, the primary goal of this analysis was to assess global differences in progenitor dynamics between sexes, rather than to resolve fine-grained spatial patterns within each tissue. For these reasons, and to increase statistical power, we pooled clones across regions when estimating mean clone sizes and performing drift model fits.

At early timepoints, most clones were composed of a single labeled cell. Over time, the average clone size increased in both the basal and intermediate compartments, while the number of surviving clones declined, as expected under homeostatic conditions (Figure 5H–K, S3D). Classical neutral drift theory predicts that clone survival should decrease approximately as ∼ 1*/t* due to stochastic competition and clonal extinction ^4^. While our data broadly follow this trend, the decline in persistence was slightly shallower than expected. Notably, the density of labeled basal cells per area remained stable between 4 and 24 weeks (Figure 6K), indicating that total labeled cell content is preserved even as clone numbers decrease.

Despite these deviations in survival dynamics, the average size of surviving clones increased linearly over time, consistent with the expectations of neutral drift. Under this model, clone size follows:

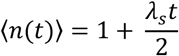

where *λ_s_* is the *effective symmetric renewal rate*, denoting the rate at which progenitor divisions generate daughters that remain within the basal compartment.

Fitting this model to the evolution of basal clone size yielded estimates of *λ_s_* = 0.47 week^−1^ for females and *λ_s_* = 0.16 week^−1^ for males. These values suggest that female Krt5^+^ progenitors undergo symmetric self-renewal more frequently than male counterparts. This sex-specific difference aligns with population-level YFP labeling (Figure 5C) and H2B-GFP dilution measurements (Figure 5O–P), both of which independently indicate faster turnover in the female urethral epithelium.

To further test model predictions, we examined the cumulative distribution of clone sizes normalized by their timepoint-specific mean (Figure S6A–B). Under neutral drift in two or three dimensions, this distribution is expected to follow:

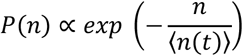

a consequence of stochastic, equipotent behavior and the progressive loss of clones due to drift ^2^. Male clone size distributions closely matched this exponential form (*R*^2^ = 0.90), while the female distributions exhibited a weaker but still qualitative fit (*R*^2^ = 0.59). The reduced fit quality in females may reflect increased heterogeneity in proliferative behavior or sampling variability. Supporting this, our Poisson analysis of H2B-GFP dilution revealed broader division rate distributions in females, suggesting greater variability in cell cycle kinetics.

Finally, the difference in magnitude between clone size scaling parameters and total division rates highlights a key conceptual distinction: clonal dynamics capture only divisions that maintain labeled progenitors within the basal compartment. In contrast, divisions that lead to differentiation or upward migration contribute to overall proliferation but not to clonal persistence. Thus, while H2B-GFP dilution measures total proliferative output, including both symmetric and asymmetric divisions, the scaling analysis provides a readout of the effective symmetric self-renewal rate.

### Krt14^+^ population-level dynamics

Following the compartment-level analysis of Krt5^+^ progenitors, we next examined the behavior of Krt14^+^ cells, a more restricted basal subpopulation. In contrast to Krt5^+^ labeling, where the fraction of labeled cells increased gradually over several weeks (Figure 5C), Krt14^+^ labeling rapidly saturated within one week and remained stable throughout the 24-week chase (Figure 6A–D, S4B).

Still, the long-term stability of labeling implies internal homeostasis — a balance between self-renewal and loss — without progressive expansion. This makes it difficult to extract quantitative turnover rates using the same modeling frameworks applied to Krt5^+^ cells. Without additional measurements (e.g., H2B-GFP dilution), we cannot resolve the division or loss rates of Krt14^+^ cells.

Despite comparable initial labeling efficiencies between sexes, we observed striking differences in epithelial tissue morphology (Figure 6C). In females, labeled cells appeared more dispersed, often as isolated single cells or loosely arranged clusters, while in males they formed compact, cohesive groups. These differences did not stem from recombination efficiency or clone size but reflected consistent divergence in spatial organization within the basal layer.

To test whether the sex-specific spatial organization of labeled Krt14^+^ cells could be explained by differences in epithelial architecture, we quantified tissue morphology using the shape index — a measure of cell packing and local tissue fluidity, defined as the ratio of cell perimeter to the square root of its area. This metric has been widely used to infer mechanical states and jamming transitions in epithelial tissues ^5–8^.

We found no significant differences in average shape index between male and female basal cells (Figure S4C–D), suggesting that individual cell geometry and local mechanical state — insofar as captured by this measure — were not detectably altered. This argues against local unjamming or cell shape changes as the cause of increased spatial dispersion in female clones.

While the exact drivers of this divergence remain unresolved, one plausible contributor is the difference in proliferation dynamics, which are more pronounced in Krt5^+^ cells and may influence spatial spread over time ^7^. Overall, these results suggest that sex-specific clonal organization emerges not from changes in local cell shape, but from broader-scale tissue behaviors that remain to be elucidated.

Together, these findings support a model in which Krt14^+^ progenitors constitute a long-lived, self-renewing subpopulation that maintains a stable presence across extended time scales. The rapid plateau and long-term persistence of labeling suggest internal replacement dynamics with minimal apparent input from upstream unlabeled populations. In parallel, the sex-specific differences in tissue morphology — especially the increased spatial dispersion of labeled cells in females — indicate that global features of epithelial organization may influence how progenitor-derived cells redistribute during homeostasis. These effects may be mediated by differences in intercellular adhesion, mechanical remodeling, or collective epithelial dynamics, and underscore the importance of considering tissue-scale architecture when interpreting clonal behavior ^9,10^.

### Krt14^+^ clonal dynamics

To complement our population-level observations, we performed clonal lineage tracing of Krt14^+^ progenitors using the *Krt14-rtTA;TetO-CRE;Rosa26^Confetti^*model. Despite labeling only basal cells at induction, Krt14^+^ clones expanded rapidly: within one week, they already comprised 4–5 cells on average and contributed visibly to intermediate and luminal layers in both sexes (Figure 6G–J, S5A,G– H). This behavior contrasts sharply with that of Krt5^+^ cells, whose labeled clones remained predominantly basal and smaller at the same time point. Crucially, while Krt14^+^ clones showed early expansion, both clone size and clone density remained largely stable from 1 to 24 weeks (Figure S5B,G–H), with no detectable clone loss. This plateau-like behavior is inconsistent with classical neutral drift, which predicts continued expansion of surviving clones and concurrent extinction of others due to stochastic competition. Instead, the Krt14^+^ data support a model in which long-lived stem cells divide asymmetrically to produce short-lived, rapidly amplifying transit-amplifying progeny—consistent with a hierarchical renewal architecture described in other stratified epithelia, including during wound healing in the skin ^11^.

This interpretation is further supported by the clear spatial separation of Confetti-labeled clones, ruling out clonal merging as a source of artificially large clones (Figure 6G–H, S5A). Moreover, both clone density and labeled basal cell density remained constant over the entire 24-week chase (Figure S5B, 6L), indicating that while individual clones neither grew substantially nor disappeared, the overall labeled population was stably maintained. These features argue against ongoing neutral competition and instead point to internally regulated output from a persistent, self-renewing Krt14^+^ pool.

Analysis of mean clone size over time in the basal and intermediate layers further supports this view (Figure 6I–J). Rather than the monotonic linear growth predicted by neutral drift, Krt14^+^ clones exhibited mild, non-monotonic fluctuations: basal clone size decreased slightly between 1 and 4 weeks before stabilizing by 24 weeks. While the precise interpretation is limited by population averaging, such dynamics are qualitatively consistent with transient early expansion followed by equilibrium—expected from a system in which fast-dividing progeny are periodically lost and replaced by a stable upstream source.

Together, these findings reveal a renewal strategy distinct from that of Krt5^+^ cells. Whereas Krt5^+^ clones undergo continuous stochastic competition, consistent with neutral drift between equipotent progenitors, Krt14^+^ cells maintain clonal output via a unidirectional, hierarchical lineage. This functional heterogeneity within the basal compartment may reflect a division of labor between stable, self-renewing stem cells and responsive amplifying progenitors, enabling the urethral epithelium to couple long-term maintenance with rapid regenerative output.

