## Supplementary material for "Two populations with different cell dynamics maintain urethral epithelial homeostasis in female and male mice": Document S4: Supplementary note on cell behaviour mathematical modelling

### Quantitative analysis of progenitor dynamics

To systematically dissect the renewal strategies of urethral progenitor populations, we implemented a quantitative framework integrating multiple complementary analyses. Specifically, we examined: (i) clonal dynamics to distinguish neutral versus hierarchical fate behavior, (ii) sex-specific differences in population-level replacement kinetics, and (iii) direct division-rate measurements using H2B-GFP label dilution.

### $Krt5^+$ population-level dynamics

To interpret the time evolution of $YFP^+$ cell fractions in *Krt5Cre-ERT2;Rosa26$^{YFP}$* mice (Figure 5C), we modeled epithelial maintenance using a simplified three-compartment framework. In this model, $Krt5^+$ basal progenitors divide and give rise to intermediate and luminal cells, which are progressively replaced over time. Let $B(t)$, $I(t)$, and $L(t)$ denote the number of labeled Basal, Intermediate, and Luminal cells, respectively. Their dynamics are governed by:

$$\frac{dB}{dt} = \lambda_b B - \lambda_i B$$

$$\frac{dI}{dt} = \lambda_i B - \lambda_l I$$

$$\frac{dL}{dt} = \lambda_l I - \lambda_d L$$

Here, $\lambda_b$ denotes the *total division rate* of basal cells, $\lambda_i$ is the *basal-to-intermediate differentiation rate*, $\lambda_l$ represents the *intermediate-to-luminal differentiation rate*, while $\lambda_d$ accounts for the *loss of luminal cells*, for instance due to cell shedding. Similar compartmental modeling framework, grounded in neutral drift theory, was previously applied by Doupé et al. to describe progenitor dynamics in the esophageal epithelium [1]. While our study includes clonal lineage tracing, this model specifically describes *population-level replacement* inferred from cumulative YFP labeling. For instance, this whole compartment dynamics is not sensitive to whether basal divisions are symmetric or asymmetric, as it represents the global fluxes between compartments.

To analyze tissue-wide labeling kinetics, we focused on the total fraction of $YFP^+$ epithelial cells over time:

$$f(t) = \frac{B(t) + I\,(t) + L\,(t)}{Total\ epithelium}$$

Under homeostatic conditions, the number of cells in each compartment remains constant, implying balanced fluxes. In particular, homeostasis requires that the rate at which basal cells divide equals the rate at which they differentiate into intermediate cells, i.e., $\lambda_b = \lambda_i$. To further simplify the system, and based on histological observations that all three compartments contain comparable numbers of cells at steady state, we assumed that:

$$B_\infty \approx I_\infty \approx L_\infty \quad \text{and} \quad \lambda_b = \lambda_i = \lambda_l = \lambda_d = \lambda$$

This approximation does not imply neutral drift — which concerns symmetric fate behavior at the clonal level [2] — but reflects a regime of uniform, balanced flux between compartments at the population level. Under these assumptions, the cumulative labeling dynamics simplify to a first-order saturation process with nonzero initial condition:

$$f(t) = f(0) + \alpha\,(1 - e^{-\lambda t})$$

where $f(0)$ is the initial labeled fraction of basal cells, and $\alpha$ captures the eventual contribution of these cells to the suprabasal layers. Since only basal cells are labeled at induction, $f(0)$ represents the baseline label and the exponential term reflects upward flux into intermediate and luminal compartments. In this framework, the long-term labeling fraction reaches:

$$f(\infty) = f(0) + \alpha$$

If compartments are of similar size and uniformly supplied by basal progenitors, we expect $\alpha \approx 2f(0)$, yielding a final labeled fraction roughly threefold higher than the initial value.

Fitting this model to the experimental data yielded a good match across all timepoints. Female urethras exhibited a faster effective turnover rate ($\lambda \approx 0.3$ week$^{-1}$), while males showed a slower rate ($\lambda \approx 0.2$ week$^{-1}$). The fitted values of $f(0)$ and $\alpha$ were both slightly higher in females, consistent with a combination of greater initial recombination efficiency and more efficient label propagation into suprabasal compartments.

Together, these results suggest that labeled $Krt5^+$ progenitors progressively populate the epithelium via steady upward transitions, and that sex-specific differences in effective replacement rates underlie the observed labeling kinetics. While the model abstracts away clonal variability and assumes homogeneous behavior, it provides a useful coarse-grained view of epithelial turnover before accounting for cell-to-cell heterogeneity.

### $Krt5^+$ division rate estimation from H2B-GFP label dilution

To independently quantify division rates of $Krt5^+$ basal progenitors, we analyzed histone H2B-GFP label dilution at 3 weeks post-labeling in *Krt5tTA;tetO;H2B-GFP* mice (Figure 5L–P, S4A). In this system, H2B-GFP is expressed constitutively in basal cells in the absence of doxycycline (DOX), resulting in a strong, uniform nuclear signal. Upon DOX administration, new H2B-GFP production ceases, and the fluorescence intensity in each cell decreases by half with every cell division. Therefore, the extent of GFP signal dilution after a fixed chase period reflects the number of divisions each cell has undergone.

Fluorescence intensities measured by flow cytometry were converted to inferred division counts, assuming ideal halving of the H2B-GFP signal per mitosis. We then constructed histograms of cell division counts and fit these with a Poisson distribution:

$$P(n) = \frac{\langle n\rangle^n e^{-\langle n\rangle}}{n!}$$

where $n$ is the number of cell divisions, and $\langle n \rangle$ is the mean number of divisions per cell over the chase period. This model assumes that divisions are stochastic and independent, with a constant probability per unit time — conditions under which division events follow a Poisson process. Such a model is commonly used to describe population-level heterogeneity in proliferation under homeostatic conditions[3].
